# Developing Foundation Models for Predicting Viral Animal Host Range in Intelligent Surveillance

**DOI:** 10.1101/2025.02.13.638012

**Authors:** Jinyuan Guo, Qian Guo, Hengchuang Yin, Yilun Han, Peter X. Geng, Jiaheng Hou, Haoyu Zhang, Jie Tan, Mo Li, Yan Zhang, Xiaoqing Jiang, Huaiqiu Zhu

**Author notes:** Corresponding author: Huaiqiu Zhu; and Xiaoqing Jiang. These authors contributed equally to this work (Jinyuan Guo, Qian Guo).

## Abstract

Emerging human infectious viruses originating from animals continue to pose a persistent threat to global public health. Understanding the host range of animal viruses is crucial for identifying potential spillover pathways and mitigating the risk of future pandemics. Here, we present VirHRanger, a prediction method that integrates foundation models trained on viral genome and protein sequences, alongside genomic and protein compositional traits, viral phylogeny, and protein-protein interactions. To systematically predict the animal host range, VirHRanger incorporates host taxonomy-aware neural networks trained on a comprehensive collection of animal-virus associations spanning mammals, birds, and arthropods. Within a dataset of 4,006 virus species spanning 99 viral families, our model achieved robust performance with a micro-averaged AUROC of 0.938 across all host categories, demonstrating its effectiveness in capturing generalizable host signals from viral genetic data. On a dataset of 315 novel viruses, which are associated with key reservoir animal hosts and insect vectors, VirHRanger notably outperformed the homology-based method, exhibiting a strong generalizability to novel viruses. Furthermore, VirHRanger identified host range variations among closely related viruses within the *Coronaviridae* family and successfully predicted the ability of SARS-CoV-2 to infect humans and other animal hosts. These findings highlight the potential of VirHRanger to transform sequencing data into timely insights for disease control during the early stages of zoonotic outbreaks.

## Introduction

In the past few decades, the majority of human infectious diseases were caused by viruses originating from wild and domesticated animals, which are known as viral zoonoses^1^. The recent COVID-19 pandemic has caused severe socioeconomic impacts and demonstrates the vulnerability of humans to zoonotic viruses^2^. Typically, viruses capable of infecting a wide range of animal hosts exhibit greater zoonotic potential^3,4^, likely due to their ability to exploit cellular machinery conserved across multiple species and their capacity to transmit through diverse animal-to-human pathways^5,6^. These generalist viruses circulating in mammals can accumulate evolutionary changes that pre-adapt them to human infection^7^. Furthermore, climate change, urbanization, and increased global connectedness facilitate the transmission of generalist viruses to previously unreachable hosts, thereby amplifying zoonotic risks and threatening endangered wildlife^3,8–11^. Therefore, preventing future disease outbreaks requires a better understanding of animal viromes and viral host ranges^1,12,13^.

Traditionally, studies on viral emergence have focused on identifying transmission routes, placing humans and reservoir hosts at opposite ends of linear chains, and linking them through intermediate hosts or vectors. The identification process, which relies on field surveillance and laboratory experiments, can take months to years and is often inconclusive, as evidenced by the ongoing debates over intermediate hosts of SARS-CoV-2^14^. However, viruses are omnipresent, and many of them frequently jump between animal hosts without causing observable symptoms. Meanwhile, viral transmissions to humans represent only the “tip of the iceberg” in the intricate network of animal-virus interactions. Therefore, several statistical models have been established to predict viral host range by incorporating phenotypic, ecological, and topological traits, which greatly expanded our knowledge of mammal-virus associations^3,15–19^. However, avian and insect hosts are often overlooked in host range predictions, even though some highly pathogenic viruses, such as West Nile virus, can infect and replicate in birds, mosquitoes, and ticks. These complex viral circulations among birds, insects, and mammals have frequently caused human disease outbreaks, making it more difficult to interrupt transmission routes^20^. Furthermore, the viral phenotypic traits used by these statistical models, such as host phylogenetic range and disease-related citations, are only available for extensively studied viruses. In addition, many traits are conserved within viral families, such as nucleic acid type and replication in the cytoplasm, while viruses of the same genus may exhibit high variability in host ranges^21–23^. Considering that most novel viruses are now discovered through metagenomic sequencing with limited phenotypic information^24–26^, models relying on sequencing data offer advantages over phenotypic trait-based models when applied to emerging viruses.

The rapid development of high-throughput sequencing technologies has revolutionized virology research^8^. The rapid accumulation of viral genomics and protein sequencing data has shed light on the molecular mechanisms of host jumping and the evolutionary history of viruses across different animal hosts^27–29^. As a result, host-specific signals have been found imprinted into viral genomes and proteins during co-speciation and host-shifting events^23,27,30^. These evolutionary signatures, such as codon preference biases of viral genomes, have been demonstrated to generalize across viral families with distant phylogenetic relationships and inform hosts of novel viruses^31–35^. Furthermore, deep learning models using Convolutional Neural Networks (CNNs) or Recurrent Neural Networks (RNNs) have been developed to predict viral or phage hosts directly from raw genome sequences^36–41^. However, most existing computational approaches rely on fully supervised learning frameworks, which are constrained by the limited availability of experimentally validated host-virus associations. Moreover, current sequencing efforts are disproportionately focused on a small portion of virus species that have caused severe socioeconomic outcomes^13,42–44^. Consequently, models trained on these overrepresented datasets are prone to sampling biases that hinder the detection of generalized host-related signals.

Recently, foundation models have shown significant promise in various biological and medical fields, such as protein structure prediction^45^, single-cell annotation^46^, and disease diagnosis^47^. These models typically incorporate hundreds of millions of parameters trained on extensive unannotated data, thereby effectively capturing complex contextual dependencies within biological data. As a result, foundation models can produce transferable representations to solve downstream tasks, particularly those with limited annotated data^48^. Recent advances in foundation models for human and prokaryotic genomes have demonstrated their strong potential to decipher biologically meaningful patterns from genetic sequences^49–51^. Moreover, conducting an independent pretraining stage on more balanced, unlabeled data enables foundation models to generate robust representations that are less influenced by the anthropocentric biases present in known animal-virus associations^52^. Benefiting from the rich sequencing data of animal viruses, which remains a largely untapped resource, we propose that foundation models can effectively extract host range information from viral genomes and proteins.

In this study, we introduce the **Virus-Host Range** predictor (VirHRanger), an integrated ensemble framework designed to systematically predict the host ranges of animal viruses. We curated a comprehensive dataset of animal-virus associations, spanning mammals, birds, mosquitoes, and ticks, establishing a basis for investigating viral host ranges and zoonoses. To decipher generalizable host signals encoded in viral genomes, we pretrained a genomic foundation model on deduplicated viral genomes and subsequently fine-tuned it on the animal-virus association dataset. Similarly, we fine-tuned a protein foundation model upon viral proteins to capture dependency patterns. We further integrated viral phylogeny, human-virus protein-protein interactions (PPIs), and sequence compositional traits from viral genomes and proteins into the predictive ensemble, considering their efficacy in prior studies. For systematic predictions of animal host range, we implemented host taxonomy-aware neural networks as classifiers. To mitigate anthropocentric sampling biases and ensure objective evaluations, we also selected a single representative genome for most virus species. As a result, the genomic and protein foundation models significantly outperformed fully supervised models based on viral phylogeny and sequence compositional traits. Our ensemble predictor, VirHRanger, achieved the best overall performance across all host categories, with a micro-averaged AUROC of 0.938. Furthermore, VirHRanger demonstrated its strong capacity to identify host range variations among closely related viruses within the *Coronaviridae* family. When applied to novel viruses from mammals and arthropods, VirHRanger accurately predicted their associated animals. Finally, VirHRanger showed promise in identifying human infecting viruses and their animal hosts during the early stages of viral emergence, as evidenced by its application to SARS-CoV-2, which can provide actionable insights for the surveillance of key animals and support the primary prevention of zoonotic pandemics.

## Results

### Comprehensive curation of animal-virus associations

Viruses are regularly exchanged among animal hosts, and some of them jump to humans from mammals, birds, or blood-feeding arthropods, causing viral zoonoses and triggering disease outbreaks. However, most host-virus resources are restricted to specific virus types or hosts^31,53^, such as coronaviruses^18^ or mammals^3^, while DNA viruses and viromes of birds or invertebrates are often neglected. Therefore, we curated a comprehensive collection of associations between viruses and animal hosts (Fig. 1a and Supplementary Table 1). Given the omnipresence of viruses in the global ecosystem, the host classifications from the traditional emergence viewpoint, such as reservoir hosts, intermediate hosts, novel hosts, and vectors, are oversimplified compared to the complicated viral exchange history among hosts^42^. Therefore, we focused on the compatibility between viruses and animal hosts and included positive associations supported by serological evidence, viral isolation, or PCR amplification. Altogether, 506,703 original records were collected from four databases, including VIRION^43^, Virus-Host Database^54^, InsectBase 2.0^55^, and Arbovirus Catalog. However, the original data are highly redundant and heavily biased to a few virus species with several social-ecological outcomes, reflecting the anthropocentric sampling bias. We reconciled virus and host taxonomy information against the unified NCBI taxonomic backbone^56^ to mitigate the bias. We harmonized the associations at the species level for both viruses and animal hosts to reduce redundancy. As a result, our collection included 4,398 animal host species and 11,320 virus species, with 28,234 unique interactions between taxonomically valid viruses and animals, representing 0.057% of all animal-host combinations. These interactions cover one-quarter (1,635 species) of mammal diversity (∼6,500 species), one-tenth (1,080 species) of birds (∼10,000 species), 6.5% (228 species) of mosquitoes (∼3,500 species), and 12.8% (90 species) of ticks (∼700 species). Furthermore, we integrated genomics into the investigation of host-virus associations by incorporated genome data from the NCBI Assembly database^57^. As a result, there are 4,006 virus species with complete genome, spanning 99 recognized viral families, while 1,039 species remain unassigned to any established viral family. In summary, our collection of host-virus associations includes mammal, bird, and arthropod hosts and covers a comprehensive spectrum of viral taxa, which provides a valuable basis for investigating viral host ranges and extracting generalizable host-related signals.

**Fig. 1.**
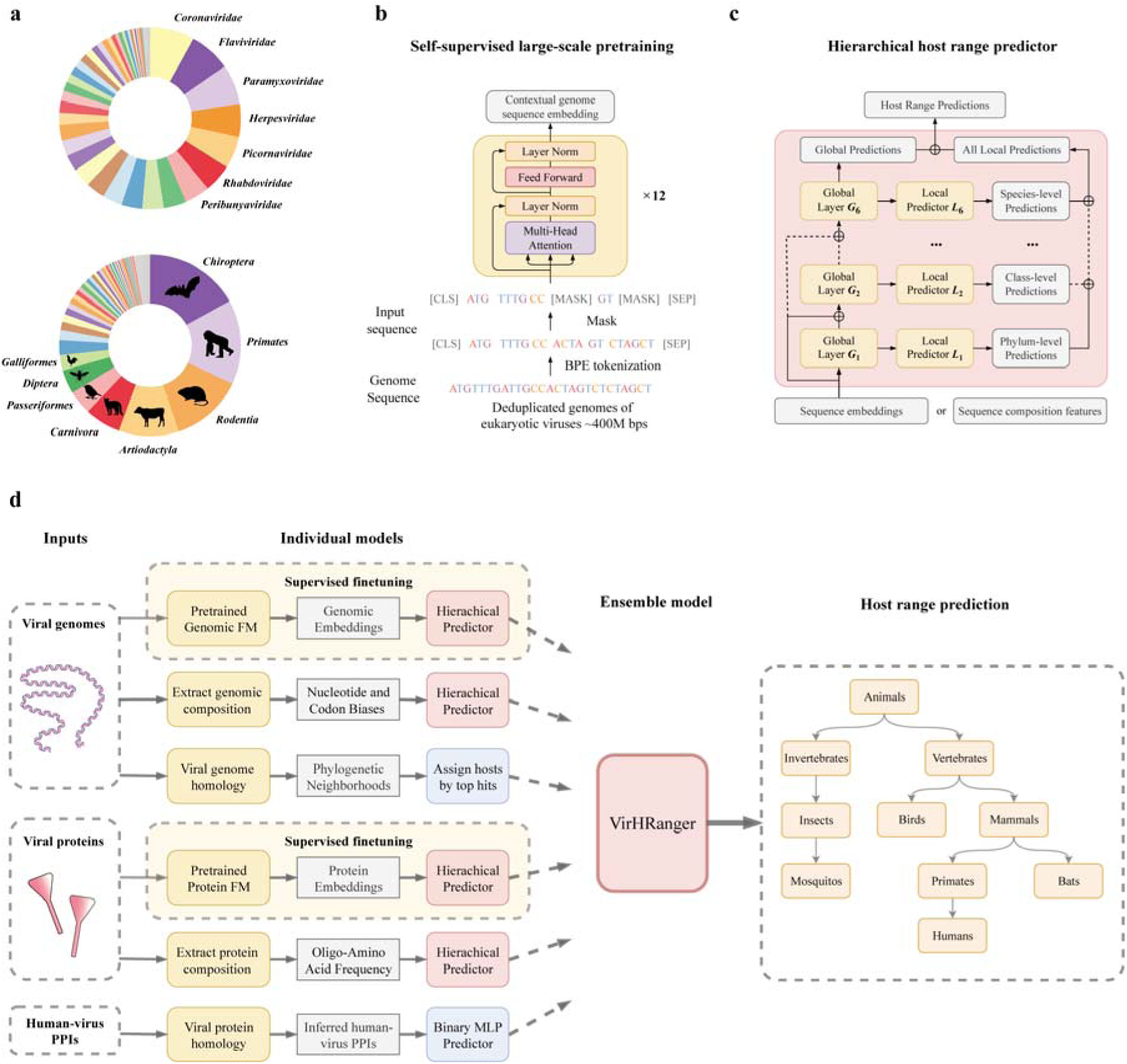
Overview of VirHRanger. **(a)** Donut charts depict the composition of the animal-virus association collection at the family level for viruses and at the order level for animal hosts. Silhouettes represent different animal host orders. **(b)** Schematic of self-supervised pretraining for the genomic foundation model. A large amount of raw viral genome sequences was tokenized, randomly masked, and fed into twelve layers of transformer encoder blocks. Encoders were trained to reconstruct the masked nucleotides based on their contextual genomic sequences. **(c)** Structure of the hierarchical host range predictor, comprising a global predictor and six local predictors corresponding to host taxonomic ranks from phylum to species levels. Input features pass through global layers and are then fed into both local and global predictors. Skip connections provide initial information to each global layer. Predictions from local predictors are concatenated and aggregated with global predictions to form the final host range predictions. **(d)** Workflow of the VirHRanger predictive ensemble. Six individual models were developed for viral host prediction: three trained on viral genomes, two on viral proteins, and one on human-virus protein-protein interactions (PPIs). The pretrained genomic and protein foundation models underwent supervised fine-tuning using host-virus association data. Contextual embeddings extracted by the foundation models were fed into hierarchical predictors for host range predictions. Additionally, genomic and protein compositional traits were utilized to train separated hierarchical predictors. The homology model infers animal hosts based on viral phylogenetic information. Furthermore, a PPI-based model was constructed to predict human viruses. Finally, the optimized six individual models were integrated into an ensemble model named VirHRanger to systematically predict host ranges of viruses at each taxonomic level. Figures were created using diagrams.net (https://www.diagrams.net).

### Architecture of VirHRanger predictive ensemble

To extract host-related signals from viral genomes, we first built a genomic foundation model for viruses using the Bidirectional Encoder Representations from Transformers architecture (BERT)^58^ (Fig. 1b). Since most existing predictors were trained either on a small number of virus species or closely related genomes^36,37^, the limited number of known host-virus associations and the uneven representation of viral families prevent these models from representing the diversity of animal viromes and extracting generalizable patterns^13^. In contrast, we pretrained a genomic foundation model on 91,169 eukaryotic viral sequences that were obtained by removing redundancy from three million viral genomes, which is more computationally efficient and may enhance the robustness of the model^59^. During the pretraining stage, viral sequences were randomly masked and reconstructed from their contextual genomic sequences, a process known as masked language modeling (MLM). We implemented the Byte Pair Encoding (BPE) algorithm^60^, which compresses viral genomes to enable longer input lengths and more effectively captures frequent oligo-nucleotide patterns. As a result, the genomic foundation model can compress raw viral genomes into contextual embeddings while preserving essential biological information.

Next, we incorporated viral protein sequences and human-virus PPI data into our predictive ensemble. Protein foundation models have been demonstrated to capture evolutionary and protein structure information and have shown promise in various applications^45,61^. Therefore, we utilized a pretrained protein foundation model named ProtBERT^61^. Considering that less than 1% of the protein sequences used to pretrain ProtBERT originated from viruses, we fine-tuned the model parameters on a large collection of viral proteins to enhance the performance of the model in the viral context. Furthermore, we incorporated host-virus PPI information, which carries essential information for viruses to enter host cells, hijack cellular machinery for replication, and escape immune responses^62–64^. Given that most documented interactions focus on human viruses, we developed a predictor for human viruses using PPI information. Specifically, we retrieved 48,643 experimentally validated human-virus PPIs from the HVIDB^65^, spanning 203 virus species. To predict hosts for new viruses, we constructed features based on homology between viral proteins and the Gene Ontology annotations of human proteins. On the other hand, we incorporated information on short linear motifs (SLiMs), an evolutionarily conserved class of protein sequences that can function as PPI interaction sites. Considering that viruses can exploit human SLiMs to their advantage through molecular mimicry, we identified potential mimicry of human SLiMs by viral proteins and built a predictor based on those SLiMs and the above-mentioned features derived from human-virus PPIs.

Using the contextual embeddings derived from viral genomes and proteins, we designed neural network classifiers to perform host range predictions. Given that currently identified host-virus associations are limited and heavily influenced by sampling biases, predicting all animal hosts at the species level remains impractical. Therefore, we selected only host categories with an adequate number of representative viruses. We started with Metazoa at the kingdom level and further subdivided a host category only when its next lower taxonomic level contained host categories with sufficient virus associations (Methods). This process resulted in 79 animal host categories spanning different taxonomic ranks, from phylum to species (Supplementary Fig. 1). Considering that many viruses can infect a wide range of animals, we framed the host range prediction as a hierarchical multi-label classification task. Specifically, we developed a taxonomy-aware classifier to systematically predict animal host ranges by employing a hierarchical multi-label classification network architecture^66^ (Fig. 1c). This architecture consists of a global predictor and six local predictors corresponding to host taxonomic ranks from phylum to species. The local predictors focus on host classes at their assigned taxonomic ranks, while the global predictor provides predictions across all host categories simultaneously. Finally, the outputs from the global and local predictors are aggregated using a weighted average. To encourage host range predictions to align with the taxonomic hierarchy, we introduced a hierarchical violation penalty in the loss function. During the training of the classifiers, we also fine-tuned the genomic and protein foundation model using low learning rates to enhance performance.

Furthermore, we developed hierarchical classifiers based on compositional traits of viral genomes and proteins, which have demonstrated effectiveness in predicting reservoir hosts and identifying viral protein functions^31,33,67^. For each viral genome, we calculated 4,229 compositional traits, including biases in codon pair usage and frequencies of dinucleotides, codons, and amino acids. These traits have been shown to capture evolutionary signals imprinted during host-virus cospeciation^31,33^. For viral proteins, we computed 11,201-dimensional compositional traits^67^, including eight protein-level features and frequencies of dipeptides, tripeptides, and k-mers features of side chain chemical groups. Moreover, viral phylogenetic relatedness is commonly regarded as a rule-of-thumb method for host prediction based on the assumption that closely related viruses tend to share similar hosts. Therefore, we developed a predictor based on the phylogenetic relatedness between viruses and used it as a baseline for benchmarking (Methods). Notably, given the limited availability of experimentally verified incompatible host-virus pairs, we assumed undocumented host-virus pairs to be negative associations during training^13^, which reflects the relative rarity of compatible interactions in nature. To improve model robustness and account for potentially unreported but compatible host-virus pairs, we assigned confidence weights to negative associations based on the viral homology (Methods). Finally, we aggregated the predictions from six individual models—comprising foundation models, compositional trait-based models, the homology-based model, and the model utilizing human-virus PPI—using an ensemble-learning strategy^68^ to form the final predictions (Fig. 1d). We suggest that the resulting ensemble model, VirHRanger, integrates contextual representations, compositional traits, and phylogenetic relatedness to achieve optimal performance in predicting viral animal host ranges.

### VirHRanger accurately predicts hosts across taxonomic ranks

To train and evaluate the predictors, we selected a single representative genome for most virus species or strains in our animal-virus association collection (Methods, Supplementary Table 2). Consequently, we obtained 25,550 interactions between animal hosts and viruses with representative genomes, which include 4,451 taxonomically resolved animal hosts (3,853 species) and 4,265 representative viral genomes (4,006 species). Cross-validation was employed for model training and evaluation. Specifically, 10% of the host-virus associations were reserved as the test set, while five-fold cross-validation was conducted on the remaining data. To enhance the robustness and generalizability of our models, we applied the CD-HIT^69^ clustering algorithm to viral genomes to minimize the sharing of homologous viruses among the training, validation, and test sets^67^. Additionally, stratified sampling was implemented to ensure balanced distributions of positive records in all host categories across the training, validation, and test sets.

Considering that most of the 79 host categories have uneven class distributions between positive and negative records, we employed metrics robust to class imbalance for model evaluations on the held-out test set. The metrics included the area under the receiver operating characteristic (AUROC), the area under the precision-recall curve (AUPRC), and the true skill statistic (TSS)^17,70–72^. To compare overall performance, we aggregated predictions across all categories using both micro-averaging and macro-averaging techniques. The ensemble model VirHRanger achieved the best performance, with a micro-averaged AUROC of 0.938 and a macro-averaged AUROC of 0.914 (Table 1). Among individual models, predictors based on genomic and protein foundation models significantly outperformed those based on sequence compositional traits in both AUROC and AUPRC (Fig. 2a, b). Additionally, foundation models demonstrated better generalizability. Among the 110 viruses in unassigned families in the held-out test set, the homology-based method performed poorly due to the absence of closely related viruses in the training data, achieving a micro-averaged AUROC of only 0.527. In contrast, the genomic and protein foundation models achieved micro-averaged AUROCs of 0.906 and 0.919, respectively, while the ensemble model VirHRanger outperformed other models with a micro-averaged AUROC of 0.941. These results indicate that the host-related signals captured by the foundation models were robust and generalizable across diverse viral families. To evaluate the sensitivity and specificity of VirHRanger, we converted prediction scores into binary classifications using class-specific thresholds. These thresholds were determined by optimizing TSS for each host category. VirHRanger achieved optimal performance, with a micro-averaged TSS of 0.744 and a macro-averaged TSS of 0.739.

**Fig. 2.**
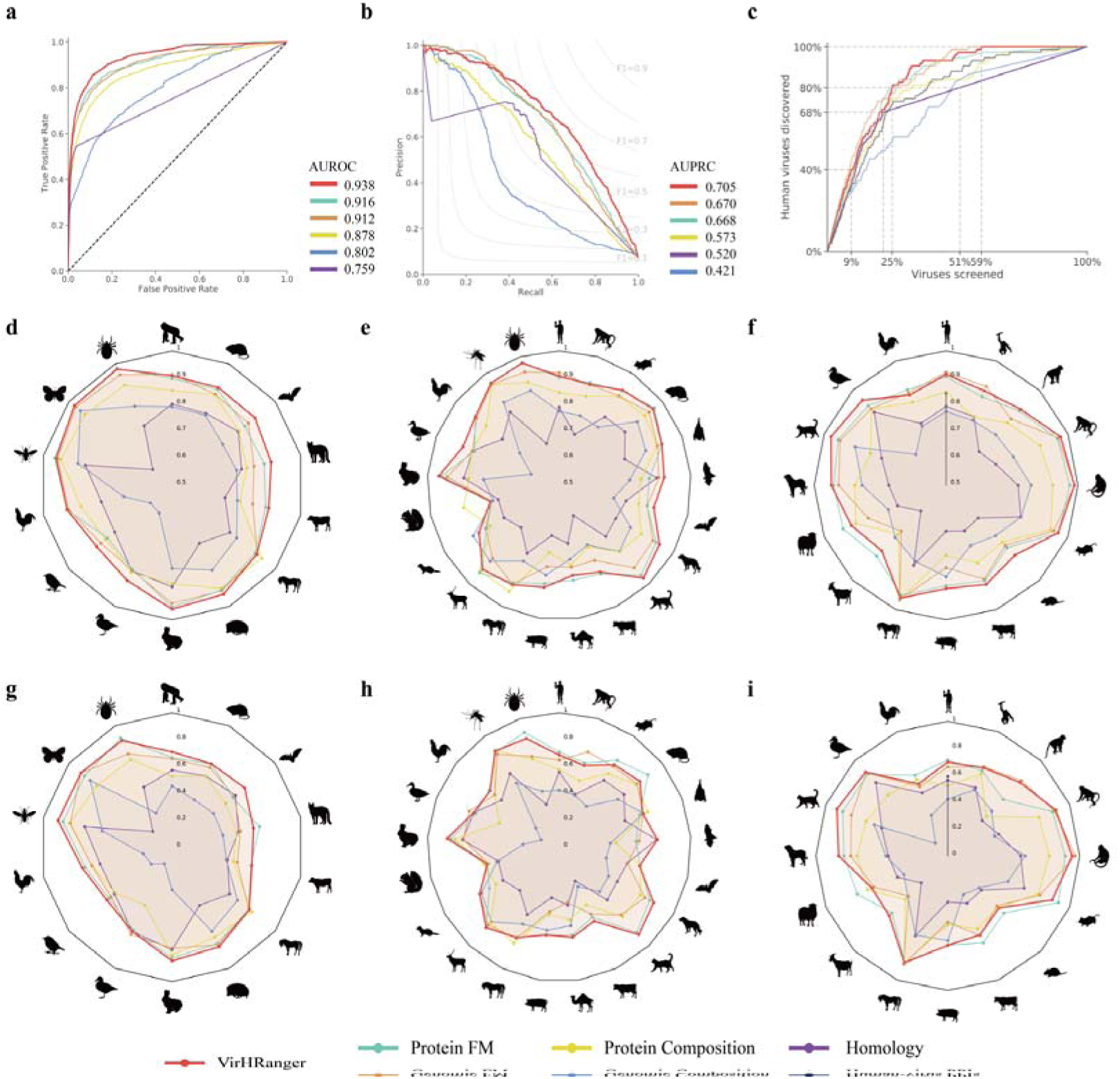
Performance of VirHRanger and individual models. **(a)** Receiver operating characteristic curves (ROC) of VirHRanger and individual models. Area under the ROC curve (AUROC) was calculated for each of the 79 host categories at different taxonomic ranks and aggregated by micro-averaging. **(b)** Precision-recall (PR) curve of VirHRanger and individual models. The area under the PR curve was calculated and aggregated by micro-averaging. **(c)** Cumulative discovery curves for human-infecting viruses of VirHRanger and other individual models. Curves show that after sorting viruses by prediction scores of different models, the proportion of animal viruses needed to be screened to discover a given percentage of known human viruses. **(d-f)** Performance of VirHRanger and individual models for specific host categories at different taxonomic ranks. AUROCs of all models in each host category at order **(d)**, family **(e)**, and species **(f)** levels. True skill statistic (TSS) of all models in each host category at order **(g)**, family **(h)**, and species **(i)** levels. Silhouettes denote different animal host categories at different taxonomic ranks. Silhouettes from PhyloPic. See Methods for image credits and licensing.

**Table 1.**
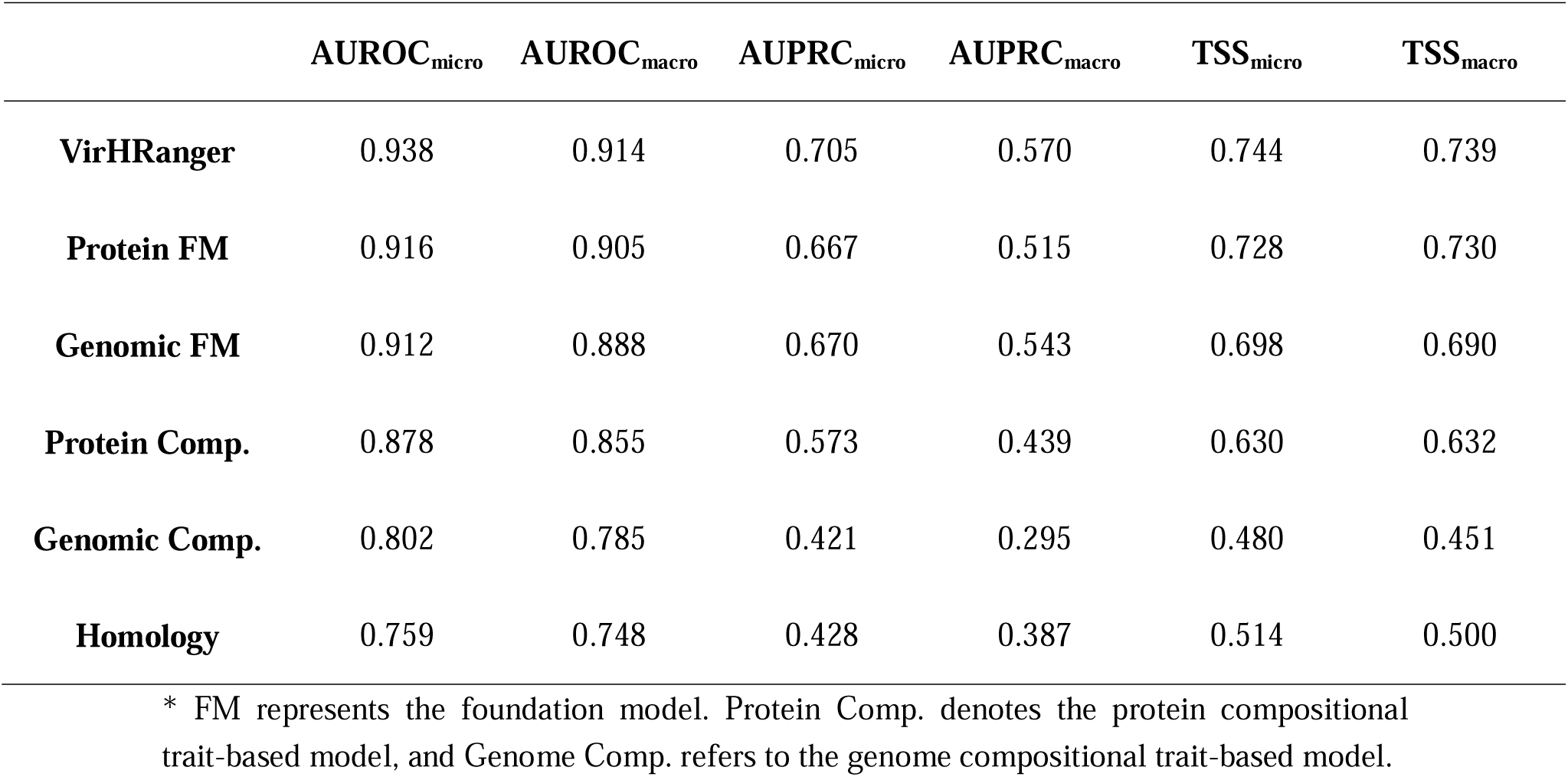
Average performance of VirHRanger across all host categories.

Notably, VirHRanger demonstrated robust predictive power at lower taxonomic ranks. Although predicting animal hosts at finer taxonomic levels is inherently challenging^35^, VirHRanger achieved comparable performance for 79 host categories across six taxonomic ranks (Supplementary Tables 3 and 4). We evaluated the performance of VirHRanger using AUROC and TSS, comparing it with individual models for each host category at the order, family, and species levels (Fig. 2d-i). VirHRanger consistently outperformed other models in most host categories, particularly in arthropod hosts. These results highlight the efficacy of our taxonomy-aware classifiers and underscore the strong predictive potential of VirHRanger for zoonotic and arthropod-borne viruses (arboviruses). It is worth noting that the human-virus PPIs-based model outperformed the genome composition- and homology-based models in AUROC when predicting human-infecting viruses (Fig. 2f and Supplementary Table 5). Despite the limited availability of known host-virus PPIs, the PPI-based model achieved AUROC comparable to the protein compositional trait-based model, demonstrating its strong potential as more PPI and protein structural information becomes available. Moreover, the PPI-based model outperformed both genomic and protein compositional trait-based models and the homology-based model in TSS (Fig. 2i and Supplementary Table 5). These findings provide a proof-of-concept for developing models by leveraging host-virus PPIs and molecular mechanisms. With rapid advancements in protein structure prediction and PPI understanding for humans and other animals, this approach could enable more accurate and insightful applications in host-virus prediction^73^.

In addition to binary host-virus association predictions, we leveraged the output confidence scores to rank the infective potential of viruses for specific hosts. This ranking approach provides a framework for guiding laboratory screening by prioritizing viruses with higher infective potential. Using VirHRanger, 40% of known human-infecting viruses could be identified by screening only the top 9.2% of viruses, while 80% could be identified by investigating the top 24.9% (Fig. 2c). Interestingly, to identify 68% of known human-infecting viruses, VirHRanger and the homology-based model demonstrated comparable efficiency when screening the top 21.5% of viruses. However, for the remaining viruses, the homology-based model relies on random guessing due to the lack of homologous signals. As a result, the homology-based model requires screening of the top 51% of viruses to achieve the same 80% identification rate, which doubles the research efforts required. To quantitatively evaluate the screening efficiency of different models across all host categories, we generated cumulative discovery curves and compared the area under the curve (AUC) across all host categories. Consequently, VirHRanger achieved the highest overall performance, with an average AUC of 0.878. In contrast, the homology-based model had an average AUC of 0.730. These findings highlight the effectiveness of VirHRanger in improving the efficiency of screening viruses with high infective potential.

### VirHRanger expands our understanding of viral host ranges

VirHRanger demonstrated strong predictive power in all host categories across different taxonomic levels, establishing a solid basis for host range prediction. Therefore, we calculated recall and TSS scores to evaluate the host range prediction for each virus in the held-out test set. We benchmarked VirHRanger against the homology-based model and found that VirHRanger consistently achieved higher performance across various taxonomic levels (Supplementary Table 6). The averaged recall and TSS scores of VirHRanger across six taxonomic levels were 0.789 and 0.605, respectively, significantly exceeding the 0.483 and 0.444 achieved by the homology-based model. Notably, VirHRanger outperformed the homology-based model by substantial margins in predicting arthropod-associated viruses. It achieved recall and TSS scores of 0.917 and 0.84, respectively, whereas the homology-based model performed poorly with scores of 0.271 and 0.241 (Supplementary Fig 2).

Next, we evaluated the predictions of VirHRanger for zoonotic and arboviruses. Here, our operational definition of a zoonotic virus is any virus species that is detected in both humans and at least one other vertebrate host without considering the transmission direction^3^. Similarly, we defined an arbovirus as any virus that is detected in both an arthropod and a vertebrate host. Following these definitions, VirHRanger successfully recalled 47 out of 50 known zoonotic viruses and identified 24 out of 27 arboviruses. Furthermore, we examined the additional zoonotic and arboviruses predicted by VirHRanger. During training, we mitigated the influence of unreported associations by assigning low confidence weights to viruses with homologous relatives infecting the host. This encouraged VirHRanger to identify undocumented but susceptible hosts for viruses. As a result, VirHRanger identified 75 additional zoonotic virus species, and 84% of them are from viral genera with known zoonotic viruses (Supplementary Table 7). Notably, we found evidence of serological or viral isolation in the literature for seven additional predictions. The family *Anelloviridae* accounted for the largest number of additional zoonotic virus predictions, with a total number of 14. Considering anelloviruses are widespread in humans and various animal species but have not been linked to any human disease, their host ranges are likely underestimated. Moreover, the update of Virus-Host DB in 2024 confirmed two of our newly predicted anelloviruses, Torque teno mini virus 11 and torque teno mini virus 12, as valid zoonotic viruses. On the other hand, VirHRanger identified 15 additional arboviruses, and we found evidence of most viruses (Supplementary Table 8). Specifically, Adana virus was recorded as having a single host, vervet monkeys, according to Virus-Host DB and VIRION database. However, literature indicated that the Adana virus can infect flies, goats, sheep, and humans^74^, and all these hosts were identified by VirHRanger. Similarly, the Kama virus was reported as tick-specific in databases, while VirHRanger predicted it had a broader host range, including birds and mammals. Recent research supported our predictions by identifying it as an atypical tick-borne bird virus^75^. Additionally, VirHRanger predicted the Southern elephant seal virus as an arbovirus, which was previously identified as the first known arbovirus of marine mammals^76^. Our findings highlighted the strong predictive power of VirHRanger in identifying potential zoonoses and arboviruses. Furthermore, we observed that many insect-virus associations in the literature have not been extracted and organized into databases, emphasizing the need for greater research focus on non-vertebrate host-virus relationships.

Given the limited knowledge of viral host ranges, we suggested that VirHRanger could identify undocumented host-virus associations, contributing to a more comprehensive understanding. Thus, we trained VirHRanger on the entire dataset and generated predictions for each virus species. Viral sharing relationships between animal host orders were analyzed separately for zoonotic and non-zoonotic viruses based on either documented (Fig. 3a) or predicted (Fig. 3b) host-virus associations. Predicted viral sharing relationships exhibited a more heterogeneous distribution compared to documented relationships, which aligned with previous findings^3,77,78^. Specifically, VirHRanger predicted a significantly higher degree of viral sharing between humans and non-human primates than reported, likely reflecting the disparity in research efforts between humans and wild primates^17^. VirHRanger also identified higher levels of viral sharing among avian orders, consistent with their phylogenetic proximity^77^. VirHRanger analyzed records of insect viruses from InsectBase 2.0, which lacked specific host information, and predicted numerous viruses in butterflies (Lepidoptera) and flies (Diptera). The analysis also revealed a high degree of viral sharing between these two host groups^79^. The high percentages of zoonotic viruses in most animal hosts highlighted the zoonotic risks and reflected the sampling biases^13,42^. Overall, the viral sharing relationships observed among mammals, birds, and insects highlight the frequent transmission of viruses between animal hosts, which poses significant zoonotic risks^42^. This underscores the urgent need for enhanced wildlife virome surveillance to control emerging zoonotic threats.

**Fig. 3.**
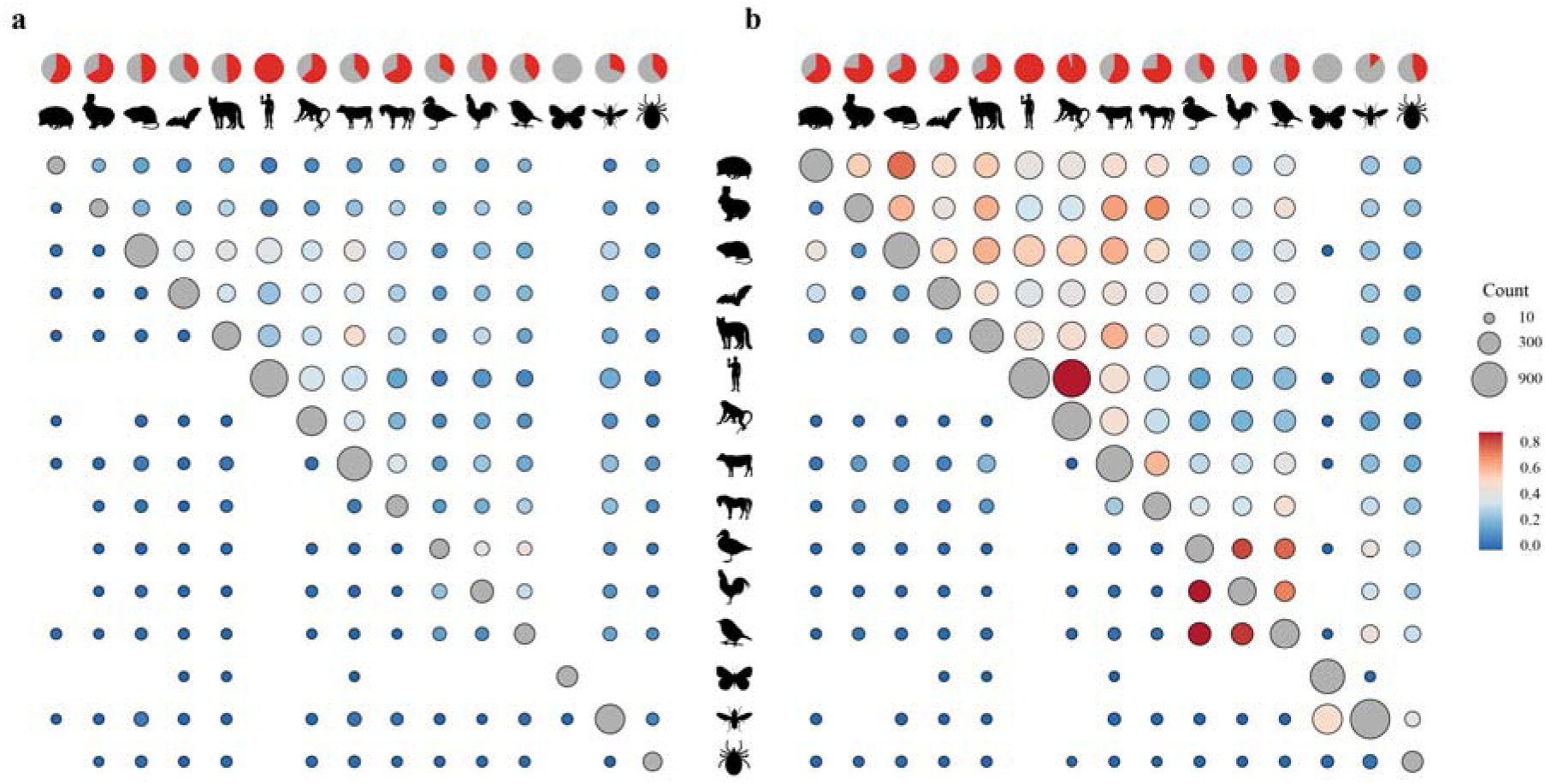
Viral sharing networks among animal hosts. Viral sharing relationships were calculated based on documented **(a)** or predicted **(b)** host-virus associations. The bubble heatmap was constructed separately for zoonotic viruses (upper triangular bubbles, excluding the diagonal) and non-zoonotic viruses (lower triangular bubbles, excluding the diagonal). The pie chart above each host indicates the percentage of zoonotic viruses in that host order, with red representing zoonotic viruses. Bubble sizes are proportional to the number of viruses shared between two host orders. Bubble color relates to the Jaccard index calculated for viral sharing. The diagonal bubbles represent the number of viruses within each host order. The primate order is split into humans and non-human primates. Silhouettes denote humans, non-human primates, and different animal host orders. Silhouettes from PhyloPic. See Methods for image credits and licensing.

### VirHRanger demonstrates generalizability to novel animal viruses

To externally validate the generalizability of VirHRanger to novel viruses, we built a validation dataset by extracting virus species that were absent from the training data in ZOVER^80^. ZOVER is a comprehensive resource for viral genomes sampled from four key animals, including bats, rodents, mosquitoes, and ticks, which are considered the most relevant animals for zoonotic and vector-borne diseases. Therefore, we used the associated animal information to examine the host predictions of VirHRanger. We removed incomplete and duplicated viral genomes for an unbiased evaluation, resulting in a dataset of 315 novel viruses (Methods and Supplementary Table 9). We evaluated the predictions of VirHRanger and the homology-based model for animal hosts across different taxonomic levels, including vertebrates, mammals, bats, rodents, arthropods, flies, mosquitoes, and ticks. As a result, VirHRanger outperformed the homology-based model across all eight host categories in AUROC, F1, and recall scores (Table 2). Specifically, for eight virus species from six genera that were absent in the training data, the homology-based model failed to predict their hosts due to a lack of phylogenetic signals. In contrast, VirHRanger successfully identified the animals associated with six virus species, demonstrating its generalizability to novel viruses. Considering that ZOVER only recorded one associated animal for each viral genome and did not account for viral sharing between animals, some additional animal hosts predicted by VirHRanger might be valid, resulting in lower but comparable precision scores for VirHRanger.

**Table 2.** External validation of VirHRanger on new viruses.

| Associated<br>Animal<br>Hosts | Host<br>Taxonomy<br>(Level) | Number<br>of<br>viruses | AUROC |  | F1 |  | Recall |  | Precision |  |
| --- | --- | --- | --- | --- | --- | --- | --- | --- | --- | --- |
|  |  |  | VHR. | Homo. | VHR. | Homo. | VHR. | Homo. | VHR. | Homo. |
| Vertebrates | Chordata<br>(Phylum) | 212 | 0.906 | 0.813 | 0.877 | 0.822 | 0.910 | 0.731 | 0.846 | 0.939 |
| Mammals | Mammalia<br>(Class) | 212 | 0.911 | 0.817 | 0.882 | 0.818 | 0.896 | 0.722 | 0.868 | 0.944 |
| Bats | Chiroptera<br>(Order) | 141 | 0.929 | 0.866 | 0.84 | 0.823 | 0.837 | 0.759 | 0.843 | 0.899 |
| Rodents | Rodentia<br>(Order) | 71 | 0.726 | 0.686 | 0.505 | 0.487 | 0.704 | 0.535 | 0.394 | 0.447 |
| Arthropods | Arthropoda<br>(Phylum) | 103 | 0.978 | 0.716 | 0.937 | 0.600 | 0.942 | 0.437 | 0.933 | 0.957 |
| Ticks | Ixodida<br>(Order) | 41 | 0.940 | 0.731 | 0.800 | 0.360 | 0.780 | 0.220 | 0.821 | 1.0 |
| Flies | Diptera<br>(Order) | 62 | 0.923 | 0.697 | 0.757 | 0.552 | 0.677 | 0.387 | 0.857 | 0.96 |
| Mosquitoes | Culicidae<br>(Family) | 62 | 0.919 | 0.700 | 0.673 | 0.558 | 0.532 | 0.387 | 0.917 | 1.0 |

\* VHR. represents the VirHRanger model and Homol. denotes the homology-based model.

Furthermore, it is important to note that in ZOVER, many viruses associated with bats or rodents were isolated from feces or digestive tracts, which raises uncertainties about whether these viruses can infect and proliferate in animal cells or are merely present temporarily due to dietary intake^81^. Similarly, viruses sampled from mosquitoes and ticks could derive from viruses parasitizing other organisms within the animals, such as bacteria or fungi, or could be ingested vertebrate viruses from blood meals^82^. Therefore, we investigated animal-virus associations that VirHRanger failed to recall by accessing their original literature (Supplementary Table 9). Among the false negative predictions, 91.3% of bat-associated and 81.0% of rodent-associated viruses were detected in digestive tracts or stools, while evidence suggests that some of them might be insect viruses^83^ (e.g., a novel *ambidensovirus*) or even bacterial viruses^84^ (e.g., four novel *picobirnaviruses*). We also found that six mosquito-associated viruses VirHRanger failed to recognize were more likely to be fungi viruses^85^, and several tick-associated viruses showed the closest relationships to viruses originating from dogs, pangolins, or cattle (Supplementary Table 9). It remains difficult to assess the infectivity of these novel viruses to their associated host, given the limited information available. However, VirHRanger can aid in screening potentially incorrect annotations for further investigation. Moreover, VirHRanger predicted that 28.6% of these viruses might infect humans, with the majority belonging to infectious disease-related viral families, including the *Nairoviridae*, *Hantaviridae*, *Bunyavirales*, and *Coronaviridae*. After excluding 17 viruses without assigned genus information, we found that 95.9% of the predicted human viruses belonged to genera known to include human-infecting viruses (Supplementary Table 9). Our findings align with estimates suggesting that a substantial proportion of viruses in wild animals have zoonotic potential^78,86^, highlighting the utility of VirHRanger for monitoring potential viral zoonotic threats in wild animals.

### VirHRanger captures variations in host range among closely related viruses

Finally, we assessed whether VirHRanger could identify different host ranges for closely related viruses. Coronaviruses (CoVs) are ideal for this purpose, as they exhibit broad and diverse host ranges across mammals and birds, with several highly pathogenic zoonotic species^87^. We predicted the host ranges for all recognized coronavirus species using their representative genomes. It is important to note that we excluded all sarbecoviruses from training data to examine if VirHRanger could effectively predict the host range of SARS-CoV-2. As a result, we constructed a phylogenetic tree to depict the phylogenetic relationships and predicted host ranges (Fig. 4 and Supplementary Table 10). The annotated dendrogram illustrated variations in host tropism among the four coronavirus genera: alpha-CoVs and beta-CoVs primarily infect mammals, whereas delta-CoVs and gamma-CoVs predominantly infect avians^88^. Exceptions to this pattern include beta-CoVs in the subgenus *Embecovirus*, which have been detected in birds^89,90^, and strains of *porcine deltacoronavirus*^91^ that were isolated from human infections. Gamma-CoVs have also been detected in marine mammals and humans. These findings highlight the broad host range of coronaviruses, which, combined with their capacity for rapid mutation and homologous recombination, continues to pose significant zoonotic risks.

**Fig. 4.**
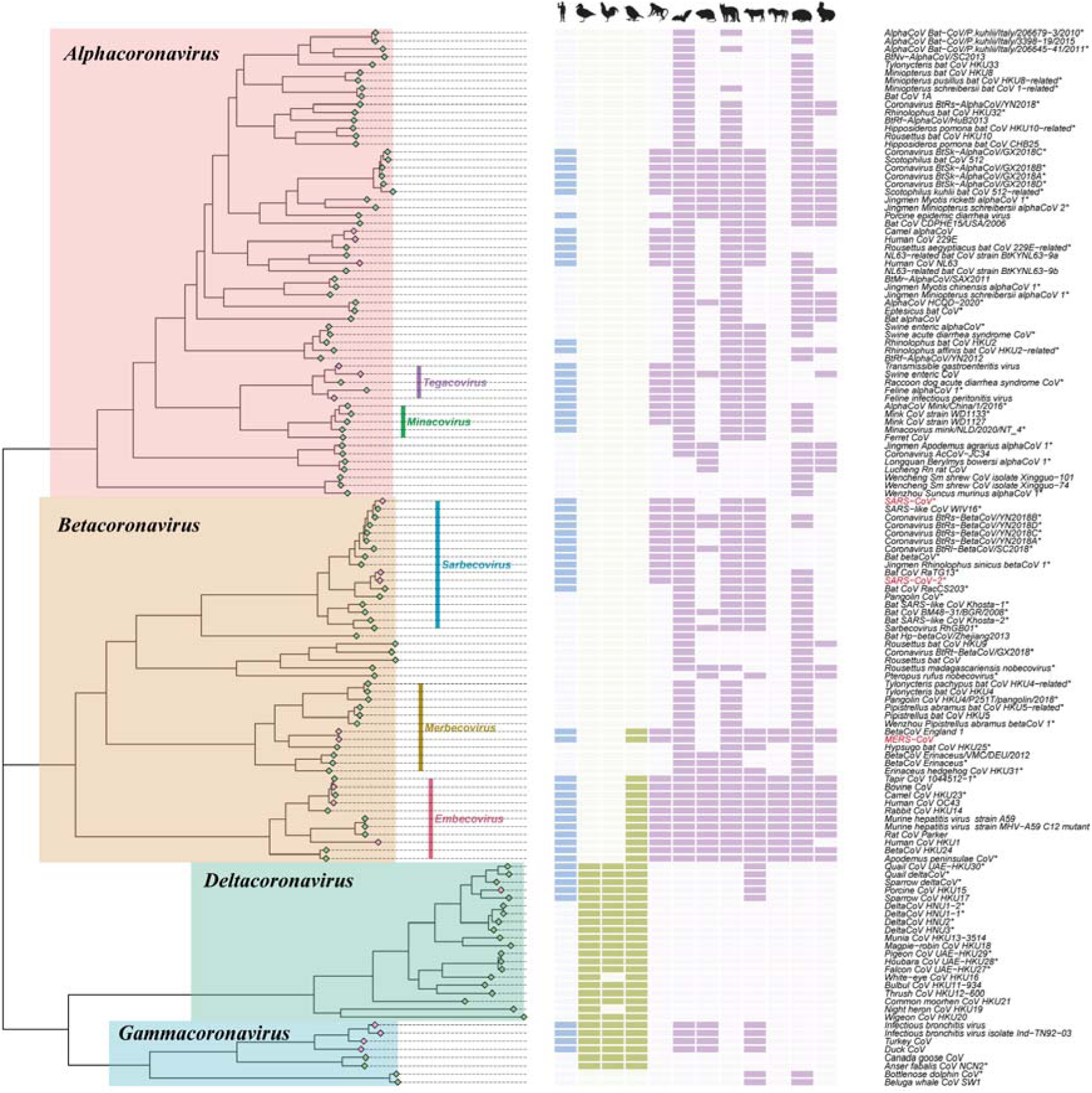
Host range predictions in coronaviruses. The Maximum Likelihood phylogenetic tree illustrates the taxonomic relationships of all recognized coronavirus species, with four colors denoting coronavirus genera. The colors at tips represent the reported virus human-infecting ability. Pink tips show known human-infecting viruses, while green tips denote those not known to infect humans. The color-coded data blocks depict animal hosts predicted by VirHRanger on order level, with different cell colors representing humans (blue), birds (green), and mammals (purple, excluding humans), and no predicted association (no color). Asterisks after species names indicate species that were not included in the training data. Three highly pathogenic species, SARS-CoV, MERS-CoV, and SARS-CoV-2, were highlighted in red. The host silhouettes, from left to right, depict humans, Passeriformes (birds), Anseriformes (waterfowl), Galliformes (landfowl), Non-human primates, Chiroptera (bats), Rodentia (rodents), Carnivora (cats, dogs, and mustelids), Artiodactyla (even-toed ungulates), Perissodactyla (odd-toed ungulates), Eulipotyphla (shrews), and Lagomorpha (rabbits). Abbreviated species names are annotated on the right-hand side. Silhouettes from PhyloPic. See Methods for image credits and licensing.

Notably, VirHRanger successfully recognized all seven commonly known human-infecting viruses in the *Coronaviridae* family, including the highly pathogenic SARS-CoV, MERS-CoV, SARS-CoV-2, and relatively mild HCoV-229E, HCoV-NL63, HCoV-OC43, and HCoV-HKU1. HKU1 and OC43 (subgenus *Embecovirus*), which have additional fusion protein hemagglutinin-esterase^92^, were predicted to have broad host ranges across mammal orders and birds. VirHRanger also predicted coronaviruses associated with cats, dogs, and minks, in subgenus *Tegacovirus* and *Minacovirus*, as potential human viruses, which aligned with clinical and computational findings^33,93^. Considering the bidirectional transmission of SARS-CoV-2 between humans and minks^38,94^, the co-circulation of alpha-CoVs and beta-CoVs on mink farms may increase recombination risks, emphasizing an urgent need for routine surveillance of novel coronaviruses in domesticated animals. Moreover, extensive studies and documented cases of SARS-CoV-2 in humans and animals have provided a more comprehensive understanding of its host range compared to other viruses. Our analysis demonstrated that VirHRanger effectively identified a broad host range, including humans, bats, cats, dogs, mustelids, and even-toed ungulates (Supplementary Fig. 3), which aligns closely with existing research findings^95,96^. Additionally, we predicted host ranges for nine SARS-CoV-2 variants of concern identified as of December 2022 to investigate their host range dynamics during evolution. The predicted host ranges were consistent across all SARS-CoV-2 lineages, suggesting that the accumulated mutations have not significantly altered their host ranges^97^. VirHRanger also predicted that bat viruses WIV16 and RaTG13 share similar host ranges with SARS-CoV-2, underscoring the potential zoonotic threats posed by viruses in the subgenus *Sarbecovirus*. Notably, since the genomes and host associations of sarbecoviruses were excluded from the training dataset, our finding highlights the predictive power of VirHRanger to effectively translate viral sequencing data into actionable insights for disease control, particularly during the early stages of virus emergence.

## Discussion

In an era of persistent threat from infectious virus emergence, the prediction and prevention of potential viral zoonoses remain unresolved challenges. The development of high-throughput sequencing and the rapid accumulation of viral sequencing data provide opportunities to establish a more complete picture of animal host-virus interactions. In this study, we curated a most comprehensive dataset of animal host-virus associations to enable robust statistical modeling of viral hosts, including not only mammals but also birds and insects. We illustrated that foundation models, pretrained on large collections of viral genomes and proteins, significantly outperformed models based on engineered compositional traits or homology in host range predictions. This finding highlights the capacity of foundation models to effectively extract generalizable host signals from virus genetic information. To incorporate homology and evolutionary insights, we integrated models of different perspectives into a powerful predictive ensemble named VirHRanger. VirHRanger identified several previously unrecognized yet valid host-virus associations and revealed the heterogeneous distribution of viral sharing between animal hosts. Furthermore, we demonstrated the generalizability of VirHRanger to novel potential zoonotic virus species and even to new genera. The ensemble model also differentiated variation in host ranges among closely related viruses. Finally, VirHRanger successfully identified the human-infecting potential and several animal hosts of SARS-CoV-2 when knowledge of SARS-related viruses was strictly excluded during model training. This highlights the utility of VirHRanger in transforming viral sequencing data into actionable insights, supporting timely disease control measures during the early stages of zoonotic outbreaks.

The limited knowledge of animal viromes remains a significant obstacle to statistical modeling. Existing experimental research is heavily biased toward pathogenic viruses of humans or a few domestic animals, leaving most animal and viral species, particularly commensal viruses, largely understudied. Even for the most extensively researched viruses, their host ranges are not fully elucidated. On the other hand, models trained on animal species with only a small number of known viruses are prone to learning sampling biases rather than biological insights. Given that very few negative host-virus relationships are documented, unreported associations are often treated as negative records during model training^13^. To address these data challenges, we applied several strategies. First, to prevent our model from being dominated by frequently sampled viruses and to enhance its generalizability to novel viruses, we conducted strict deduplication, selecting a single representative genome for most virus species. Furthermore, our genomic and protein foundation models were pretrained on balanced, unlabeled sequence datasets. This allowed the models to extract host-related signals from a limited set of deduplicated host-virus associations, which is challenging for fully supervised deep-learning models. Second, we made a trade-off between taxonomic resolution and reliability by only predicting for host categories with a sufficient number of records, setting the threshold at 80 representative viral genomes. For host categories with sparse data, VirHRanger provided host predictions at higher taxonomic levels. We also implemented a taxonomy-aware predictor, incorporating host taxonomic information to guide the information flow within the model. Our framework is flexible and can accommodate more hosts as more host-virus associations become available or when a looser threshold is applied. Third, to alleviate the impact of unobserved but valid host-virus associations, we assigned low confidence weights to negative records when viruses had homologous relatives infecting the same host. This is a common strategy to account for unknown positive records^98^. We demonstrated its effectiveness in predicting zoonotic and arboviruses, as VirHRanger identified multiple valid host-virus associations that were not included in existing databases.

Another limitation lies in the quality of data on known host-virus associations. We have curated a comprehensive collection of animal-virus associations, including a significant focus on birds and arthropod viruses. However, these records were derived from various experimental assays, such as PCR amplification, serology, or viral isolation, that vary in reliability^13,43,99^. Although we excluded experimental and cell line infections when compiling the dataset, some associations from source databases may still rely on less reliable evidence or unpublished data, potentially containing errors that are difficult to identify. Additionally, we included both pathogenic and commensal host-virus associations; however, the epidemiological roles of hosts, such as reservoir, vector, or dead-end hosts, were not considered. Therefore, VirHRanger is not designed to predict the pathogenicity of the viruses or the roles of hosts in transmission cycles. However, VirHRanger demonstrates the strong predictive power of foundation models for virus phenotypes and highlights the utility of foundation models in forecasting viral pathogenicity. Furthermore, VirHRanger does not incorporate data on host geographical distributions and, therefore, is not suitable for predicting viral transmission opportunities between hosts^11^. Given the limited availability of animal species distribution data, we acknowledge that predicting transmission opportunities remains an important but challenging task, particularly in the era of global warming, increased international travel, and urbanization.

Our findings demonstrate the principle that viral host specificity is encoded in viral genomes and proteins. Viral genetic information reflects an interweaving of virus-host co-divergence and cross-species transmission^100^, which also encodes the potential to infect new hosts. Therefore, VirHRanger was developed to systematically predict viral host ranges across different taxonomic levels rather than focusing solely on humans or reservoir hosts. We demonstrated that VirHRanger successfully captured the heterogeneity in host-virus sharing relationships. Furthermore, considering that host-virus interactions are predominantly mediated by protein bindings, we went beyond viral sequence information and designed a model integrating human-virus PPI information. Despite known PPIs representing only a minuscule fraction of the complex human-virus PPI network, our model outperformed both the homology-based model and sequence compositional models, validating the integration of molecular-level information as a proof-of-concept. Although accurately modeling the folding and clipping of viral polyproteins remains challenging, the rapid development of advanced deep learning methods, such as Alphafold3^73^, demonstrates potential in elucidating the molecular mechanisms underpinning host-virus associations.

Leveraging contextual information extracted by genomic and protein foundation models, VirHRanger effectively captured generalizable host-related signals and accurately predicted the animal host range. Recently, more advanced foundation model architectures have been developed, significantly expanding their perception fields to 1 million base pairs^50^. Furthermore, beyond guiding zoonotic virus screening, the fast and resource-efficient fine-tuning of foundation models facilitates on-the-fly training of the predictor using the feedback from field studies. Together with high-throughput metagenomic sequencing and large-scale serological profiling^101^, pretrained foundation models show great promise toward decoding the language of viral host specificity and cross-species transmission.

## Methods

### Datasets

#### Curation of animal-virus association dataset

The transmission cycles of zoonotic viruses typically involve mammals, birds, mosquitoes, and ticks. To capture the breadth of these interactions, we curated a comprehensive dataset of associations between viruses and animal hosts, including both vertebrates and arthropods. The original dataset was sourced from four databases: VIRION^43^, Virus-Host Database^54^, Arbovirus Catalog^102^, and InsectBase 2.0^55^, which together provided 506,703 records. We retrieved the data from these four sources in January 2022. The original host-virus association records were heavily biased toward well-documented virus species. To reconcile inconsistencies in virus and host taxonomy, we standardized the data by cross-referencing the taxonomy information against the NCBI taxonomic backbone^56^ using the package *taxize*^103^ in R. This allowed us to normalize the data at the species level, reducing redundancy and creating a unified dataset. As a result, the final dataset includes 4,398 animal host species and 11,320 virus species, with a total of 28,234 unique interactions between taxonomically valid viruses and animals (Supplementary Table 1). The majority of these interactions (82.6%) came from VIRION, followed by VHDB (15.7%), the Arbovirus Catalog (8.8%), and InsectBase 2.0 (5.1%). Specifically, 11.5% of these interactions were documented in more than one database.

#### Selection of representative viral genomes

We accessed the NCBI Assembly database^57^ on June 14, 2022, and retrieved all virus genome assemblies. Only records with complete genomes were retained to maintain consistency. To avoid overestimating model performance, we selected a single representative genome from each virus species, particularly to prevent the inclusion of nearly identical genomes in both model training and validation sets. Reference sequences from RefSeq^104^ were prioritized when available; otherwise, the longest genome in GenBank^105^ was selected. Exceptions were made for a small subset of extensively studied virus species that have multiple reference genomes. For instance, Influenza A virus had seven representative genomes, while Norwalk viruses had twelve. As a result, 3,839 virus species were represented by a single genome, while 167 species had more than one representative genome. This step reduces redundancy and is crucial for enhancing the model’s generalizability, especially for newly discovered viruses that may lack homologous reference genomes. Additionally, subviral agents such as satellites and viroids were excluded from the dataset. When combined with the animal-virus association dataset, we obtained 25,550 interactions between animal hosts and viruses with representative genomes (Supplementary Table 2). This dataset includes 4,451 taxonomically resolved animal hosts (3,853 species) and 4,265 representative viral genomes (4,006 species).

#### Pretraining data for genomic foundation model

For the pretraining of our genomic foundation model, we aimed to include a diverse set of viral genomes to improve generalizability. However, the currently available viral genomic sequences are highly redundant, with some species disproportionately represented. For example, SARS-CoV-2 sequences (∼9 million) constitute 69% of the total viral sequences (∼13 million) in the NCBI virus database^106^. Since non-redundant training data can improve the robustness and generalizability of the foundation model, we accessed Virosaurus^107^, a database that provides unbiased and deduplicated eukaryotic viral genomes. Virosaurus clusters viral genome sequences at a 98% identity threshold using the CD-HIT tool^69^. We used the version of Virosaurus updated in April 2020, which contains 91,169 eukaryotic viral sequences.

Since genomes of RNA and DNA viruses were stored as standard nucleotide bases (A, T, C, G), we replaced all ambiguous nucleotides with “N”. As many viral genomes exceed the input length limitation of our model architecture, we adopted a modified version of the DNABERT^49^ preprocessing method. Specifically, we processed viral genomes in three ways, and each chunk was treated as an independent sequence during pretraining:

1. Splitting long genomes: Long viral genomes were divided into non-overlapping chunks of 3,000 base pairs. Viral genomes shorter than this threshold were directly included.
2. Random sampling of long genomes: For genomes exceeding 3,000 base pairs, we randomly sampled chunks. With a 50% probability, the chunk size was 3,000 base pairs; for the remaining 50%, the chunk size was randomly chosen from a range of 10 to 3,000 base pairs.
3. Random sampling of short genomes: For genomes shorter than 3,000 base pairs, we randomly sampled chunks. The chunk size was randomly chosen from a range of 10 base pairs to the full length of the genome.

### Genomic foundation model for animal viruses

Foundation models were primarily developed for human language processing and have since achieved state-of-the-art performance across a variety of bioinformatics and medical tasks. By definition, a foundation model is pretrained on an extensive amount of data and can be accommodated to various downstream tasks, especially those with a limited amount of labeled data^108^. Masked language modeling is commonly employed during the pretraining stage. MLM task is to predict masked portions of sequences based on the surrounding context^58^, which leverages large unannotated datasets and allows the model to learn underlying dependency patterns. Most existing foundation models are based on the transformer architecture, which relies on a self-attention mechanism to transform input data into useful embeddings. A widely used architecture within this framework is BERT, which has demonstrated state-of-the-art performance in various machine learning tasks.

#### Architecture

Our genomic foundation model consists of twelve transformer encoder layers, each containing a multi-head self-attention layer and a feed-forward layer. The input size for each encoder is 512 tokens, and both the self-attention and feed-forward layers have a size of 3,072. Each self-attention layer contains twelve attention heads. The model begins with an embedding layer that transforms the input sequence of tokens into embeddings. Each token is mapped to a 768-dimensional vector. Positional embeddings are added to capture the sequential order of tokens. The model hyperparameters are as follows: a Gaussian Error Linear Unit (GELU)^109^ is used as the nonlinear activation function; the dropout probability for all hidden layers is set to 0.1; the dropout rate for attention probabilities is also 0.1; and the epsilon value for layer normalization is 1 × 10^□^¹². Model development and training were conducted using *PyTorch*, with the Huggingface *Transformers* library ^110^ facilitating data loading, configuration setting, and model training.

#### Tokenization of input genome sequences

To model nucleotide sequences of viral genomes, we built a BPE table^60^ for efficient compression of sequence length and pattern extraction. The vocabulary size was set to 32,000 tokens, and the *SentencePiece* tokenization algorithm^111^ was applied to unannotated viral genomes. This transformed sequences of nucleotides into sequences of oligonucleotides (tokens). Each token represents, on average, 10.24 base pairs in genomic sequences. Additionally, we added special tokens into the vocabulary to represent the beginning of sequence ([CLS]), end of sequence ([PAD]), and masking part ([MASK]). We used the same BPE table during pretraining and downstream applications. The same BPE table was used during both pretraining and downstream applications. Given that the computational cost of training BERT-like models grows quadratically with input length^112^, the BPE tokenization alleviates the length bottleneck and enables the model to capture dependency patterns over longer sequences.

#### Pretraining

The model was pretrained according to the BERT methodology^58^. Processed genomic sequences were tokenized using the BPE table, with special tokens [CLS] and [SEP] added at the beginning and end of each sequence. Following the observations from BERT and RoBERTa, we trained the model using the MLM task, wherein 15% of the tokens in each sequence were randomly masked. To minimize the mismatch between pretraining and fine-tuning, we employed the following masking strategy: 80% of the masked tokens were replaced by the special token [MASK], 10% were replaced by random tokens, and 10% of the tokens were left unchanged. Self-supervised training was then performed, where the model learned to predict the masked tokens based on their surrounding context. The batch size was set to 2,000, and the cross-entropy loss was computed and summed for each batch. To improve predictive power, we employed a two-phase training approach: the model was trained for 100 epochs with 15% of tokens masked and then trained for 50 epochs with 20% of tokens masked. We applied a warm-up strategy for both training stages. Specifically, the learning rate linearly increased to its peak during the first 10% of total steps and then decreased linearly to zero. The peak learning rate for both phases was set to 6×10^−4^. The Adam optimizer was used with the following parameters: *β*_1_ = 0.9, *β*_2_ = 0.98, *∈ =* 1×10^−6^, and weight decay to 0.01. All experiments were conducted on two NVIDIA A100 GPUs, with the two-stage pretraining process taking approximately 21 days to complete.

#### Fine-tuning

For fine-tuning, we split the representative viral genomes into overlapping subsequences of 3,000 base pairs, which were tokenized using the same BPE table. Special tokens [CLS] and [SEP] were added to the beginning and end of each subsequence. The input sequence of amino acids was transformed into embedding by passing it through the encoder layers. Following observations from RoBERTa^113^, the embedding vector corresponding to the [CLS] token effectively aggregates information from the entire subsequence. Hence, this vector was passed into classifiers for supervised learning. The number of training epochs was set to 50, and the batch size was set to 256. To prevent instability during training, we applied the same warm-up strategy as during pretraining. The peak learning rate for the genomic foundation model during fine-tuning was set to 1×10^−5^, which was smaller than the learning rate used during pretraining. The peak learning rate for the classifier was set to 1×10^−3^, with a dropout rate of 0.5. The same optimizer settings as the pretraining stage were used. To avoid overfitting, we employed an early stopping strategy.

### Protein foundation model for animal viruses

We downloaded the pretrained ProtBERT model^61^ from the Huggingface repository. The ProtBERT model consists of 30 layers of transformer encoders and was trained on 216 million protein sequences from UniRef100^114^. Since viral proteins are underrepresented in UniRef100, we fine-tuned the ProtBERT model on viral proteins with a small learning rate. For each representative viral genome in our dataset, we obtained all corresponding protein sequences from RefSeq and GenBank. The protein sequences were then truncated into overlapping subsequences with a maximum length of 510 amino acids. Special tokens, [CLS] and [PAD], were added at the start and end of each subsequence. We replaced the final layer of the model with a hierarchical classifier. The input amino acid sequence is transformed into embeddings by the encoder layers. The embedding generated for the [CLS] token is then fed into the classifier for supervised learning. The hyperparameter settings used were the same as those in the fine-tuning stage of our genomic foundation model.

### Gene compositional traits-based model

Previous research^31,33^ has demonstrated that virus genomic compositional biases can inform virus evolution history and their specificity to animal hosts. Hence, we incorporated these expert-designed genomic compositional traits into our predictive framework. We calculated these traits for each viral genome, and for segmented viruses, we calculated features for each segment separately. Annotations for coding regions were acquired from RefSeq and GenBank. The genomic compositional traits included in our analysis are codon biases, amino acid biases, dinucleotide biases, and codon pair biases:

#### Codon biases

Codon biases were calculated by:

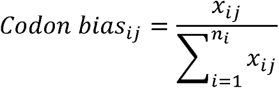

Where *x_ij_* is the number of occurrences of the *j*^th^ codon for the *i*^th^ amino acid, and *n_i_* is the number of synonymous codons that encode for the *i*^th^ amino acid. A stop codon is considered an amino acid here. There are 64 codon bias traits.

#### Amino acid biases

Amino acid biases represent the observed frequency of specific amino acids. A stop codon is treated as an amino acid. As such, there are 21 amino acid bias traits.

#### Dinucleotide biases

Dinucleotide biases were calculated by:

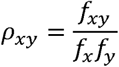

Where *f_xy_* denotes the observed frequency of the dinucleotide *XY*, and *f_x_* and *f_y_* are the observed frequencies of nucleotides *X* and *Y*, respectively^115^. Given that dinucleotide biases at “bridge” positions exert the strongest effects on viral fitness, we also calculated 16-dimensional dinucleotide biases for both “bridge” and “non-bridge” codon positions separately. This yields 48 dinucleotide bias traits.

#### Codon pair biases

Codon pair biases (64 × 64) were calculated for each possible codon pair, including stop codons. The bias was defined as the natural logarithm of the ratio between the observed number of a specific codon pair and its expected frequency over coding regions^116^.

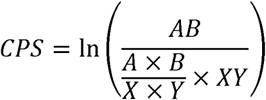

Where A and B represent the observed number of *A* and *B*, *X* and *Y* represent the observed number of corresponding amino acids, *AB* is the observed number of codon pair *AB*, and *XY* is the observed number of corresponding amino acids. The CPS scores show if a given codon pair is over-represented or under-represented. The CPS scores indicate whether a codon pair is overrepresented or underrepresented. If an amino acid pair is not observed, all corresponding codon pairs are assigned the average CPS score for the entire sequence. When the amino acid pair is observed but the codon pairs are absent, the CPS is set to -10, indicating extreme underrepresentation. This results in 4,096 codon pair bias traits.

In total, we calculated 4,229 genomic compositional traits. We then trained a hierarchical classifier based on these traits. The model was trained for 100 epochs with an early stopping strategy to avoid overfitting. A grid search was used to determine the best set of hyperparameters, which identified the optimal batch size as 16, the learning rate as 1×10^−4^, and the dropout rate as 0.1. The optimizer and other training settings were consistent with those used in the genomic foundation model.

### Protein compositional traits-based model

Researchers have demonstrated that viral protein functions can be inferred from their amino acid compositional traits^67^. We incorporated protein compositional traits, as defined in PhANNs^67^, into our predictive framework. These traits include amino acid biases, side chain chemical group biases, and eight protein-level features.

#### Amino acids biases

Amino acid biases refer to the frequency of dipeptides (400 combinations) and tripeptides (8,000 combinations) in the protein sequence.

#### Side chain chemical group biases

Amino acids can be classified into seven groups based on their side-chain chemical properties. The frequency of side chain two-mers (49 combinations), three-mers (343 combinations), and four-mers (2,401 combinations) are calculated to capture the chemical group biases in the sequence.

#### Protein-level features

We applied the *ProteinAnalysis* function from the *Biopython* package^117^ to calculate eight distinct protein-level features for each viral protein sequence, including the isoelectric point, instability index, ORF length, aromaticity, two distinct molar extinction coefficient, hydrophobicity, GRAVY index, and molecular weight.

The total number of protein compositional traits was 11,201. These traits were extracted from each viral protein and used to train a hierarchical classifier. The training settings were consistent with those used in the genomic foundation model. A grid search was used to determine the best set of hyperparameters, which identified the optimal batch size as 32, learning rate as 1×10^−3^, and dropout rate as 0.1.

### Homology-based model

The animal-virus association collection includes highly divergent viruses from various viral families, making sequence alignment difficult due to the lack of homologous genes. To address this, we adapted the phylogenetic neighborhood method^31^ defined by Babayan et al. Specifically, we performed a BLAST search^118^ for each query virus in the test set against the training set to identify its closest relatives. We used the following BLAST parameters: maximum number of high-scoring pairs = 1, reward = 2, word size = 8, gap open = 2, gap extend = 2, e-value threshold < 0.001, percent identity > 30, percent query coverage > 60. We selected the top five hits based on e-values. The host range information from the top viral genomic hits was then assigned to the query virus, with a confidence score calculated as the average of pairwise genetic identities between the query and the top five hits.

### Human-virus PPI-based model

#### Experimentally validated PPIs

We retrieved all experimentally validated human-virus PPIs from the HVIDB^65^, which includes 48,643 PPIs across 203 virus species from 35 viral families. These interactions involve 7,796 human proteins and 2,104 viral proteins. Although viral proteins account for only ∼3.5% of the proteins in our animal-virus collection, homologous signals exist in the remaining proteins. Thus, we used the 2,104 viral proteins as bridges to propagate known human-virus PPIs to new viral proteins by homology. Specifically, we set these 2,104 viral proteins as targets and performed protein-protein BLAST for all 60,636 viral proteins using default parameters with an e-value threshold of 0.001. Only the top hit for each viral protein was retained. As a result, each viral protein was represented by a 7,796-dimensional feature vector defined by bit scores. To reduce redundancy and noise while preserving functional relationships, we employed Gene Ontology (GO) annotations^119^ to compress the feature vector into a lower-dimensional representation (2,364 dimensions). Features were summed under the same GO term, describing protein functions, locations, and biological processes.

#### Computationally inferred SLiMs

SLiMs are short protein functional modules important in PPIs and post-translational modifications. SLiMs typically consist of three to ten consecutive amino acids, are linearly organized in 3D structure, and frequently occur in intrinsically disordered regions of proteins. SLiMs are critical in cellular processes such as signaling, localization, degradation, and cleavage. Viruses also exploit SLiMs through molecular mimicry. We downloaded all SLiM classes from the ELM database (2022 release)^120^. These motifs include eleven cleavage-related motifs, 26 degradation-related motifs, 38 docking-related motifs, 181 ligand-related motifs, 37 modification-related motifs, and 26 target-related motifs. We used the *SLiMProb* function from the *SLiMSuite* Python package^121^, with the default parameters. The disorder regions were predicted using IUPred^122^. We extracted the number of occurrences and the difference between observed and expected occurrences for each SLiM motif class, removing any classes with no hits. This resulted in 263 features for motif occurrence and 263 features for motif occurrence difference.

#### Model

Given the limited knowledge of host-virus PPIs and SLiMs for species other than humans, we constructed a multi-layer perceptron to identify human-infecting viruses, which comprises an input projection layer, three hidden layers with ReLU activation, and dropout for regularization. We added confidence weights to binary cross-entropy loss to address class imbalance (details described in section Algorithm optimization and validation). We followed the same dataset partitions with other models and trained the model based on features derived from human-virus PPIs and SLiMs. We trained models for 100 epochs and employed an early stopping strategy to avoid overfitting. A grid search was used to determine the best set of hyperparameters, which identified the optimal batch size as 64, learning rate as 1×10^−5^, and dropout rate as 0.1. Experiment results demonstrated that incorporating GO information enhanced model performance (Supplementary Table 11).

### Taxonomy-aware classifier

#### Hierarchical host categories

Known host-virus associations are disproportionally concentrated on humans and several domesticated animal species, while the virome of most animal species remains underdocumented and subject to sampling biases. To account for this, we framed the host range prediction problem as a hierarchical multi-label classification task by defining hierarchical host categories. Starting at the kingdom level, we subdivided a host category only if its next lower taxonomic level included host categories with a sufficient number of associations with viruses. Host categories with at least 80 known associations with representative viruses were included in the host pool and were further subdivided until the condition mentioned above was not met. Our design was inspired by Babayan’s eleven host categories at the superorder, order, suborder, or family level^31^. We expanded the scope by incorporating host categories from the phylum to the species level, which helps to study the virus-host range systematically. Ultimately, 79 host categories spanning different taxonomic levels were retained, covering animals of economic importance as well as key hosts and vectors for zoonotic viruses (Supplementary Fig. 1).

#### Architecture of the classifiers

Considering the hierarchical nature of host taxonomy, we adapted the structure of a state-of-the-art neural network architecture for hierarchical multi-label classification, known as HMCN-F^66^. Our classifier consists of six local predictors corresponding to host taxonomic ranks from phylum to species and a global predictor. The global and local predictions are aggregated into a final prediction using weighted averaging. The architecture includes six global hidden layers and six local hidden layers. The hidden size is set to 768 for global layers and 384 for local layers. Hidden layers are fully connected with ReLU activation. Batch normalization and dropout were implemented for regularization. There are residual connections between the global hidden layers. Since animal hosts are not mutually exclusive, we implemented the 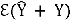 and sigmoid activation for both global and local predictors. The global loss 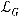 and local loss 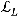 are calculated as follows:

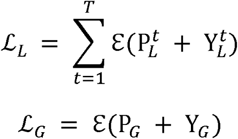

where 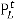 denotes the local prediction at host taxonomic level *t*, and P*_G_* denotes the global prediction. Additionally, we employed a penalty loss term^66^ to avoid hierarchy violations, i.e., when the predicted probability of a host category exceeds that of its parent category:

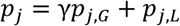

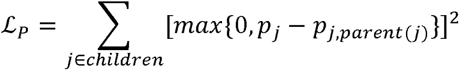

when p*_j_* is the weighted average of predicted probability of the *j*-th host category. The final classification loss 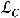 is defined as:

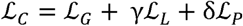

Wher*e* γ and δ are regularization parameters and are set to 1 and 0.1, respectively, in all experiments. After evaluating the final predictions, we found a small proportion of hierarchy violations and conducted post-processing to remove these violations. Finally, the predicted probability is calculated for each host category for each subsequence. The predictions of subsequences are then aggregated into a final sequence prediction by averaging.

### Algorithm optimization and validation

#### Data preparation for training and cross-validation

The dataset of associations between animals and viruses with representative genomes enabled the supervised training of our base models and meta-models. Given the limited availability of experimentally verified negative host-virus pairs, we assumed that undocumented host-virus pairs represent negative associations in supervised training. This reflects the relative rarity of compatible interactions in nature. Based on the hierarchical host categories, each representative virus genome was assigned a set of 79 host labels, indicating the presence or absence of an association between the virus and the host category. Consequently, we obtained a dataset of 4,265 representative viral genomes with host labels. We set aside 10% of viral genomes as a held-out test set and conducted five-fold cross-validation on the remaining 90% of the viral genomes for robust evaluation. To capture generalizable signals and avoid information leakage from closely related viruses present in both the training and test sets, we performed sequence clustering. Specifically, we used CD-HIT-EST to cluster viral genomes with an identity threshold of 80%, which is recommended to prevent over-clustering. This resulted in 3,995 clusters, and we then employed the *MultilabelStratifiedKFold* function from the *iterative-stratification* Python package^123^ to ensure a balanced distribution of host labels. A total of 60,636 viral proteins were annotated in the representative genomes. We followed the aforementioned data partitions, ensuring proteins from the same genome were placed in the same partition. We trained and optimized models on cross-validation sets by performing grid searches over a wide range of hyperparameters. Finally, we evaluated model performance and generalization capabilities on the independent test set.

#### Evaluation metrics

We employed metrics robust to class imbalance for model benchmarking, including AUROC^70^, AUPRC^71^, and TSS^72^. TSS accounts for both sensitivity and specificity and is defined as:

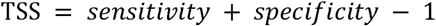

To evaluate the overall performance of the models across host categories, we employed both micro-averaging and macro-averaging. Micro-averaging aggregates all predictions and true labels across host categories, while macro-averaging calculates performance metrics independently for each host category and then computes their mean. We also calculated the F1, recall, and precision scores to compare the model performance.

#### Confidence weights for negative associations

Since negative host-virus pairs in our dataset may contain unreported but compatible associations, we assigned a confidence weight to each negative association based on viral homology during training^124,125^. Specifically, for each host category, we obtained a list of viruses with positive associations and assigned a similarity weight ***w****_sim_* based on the top hit in BLAST results:

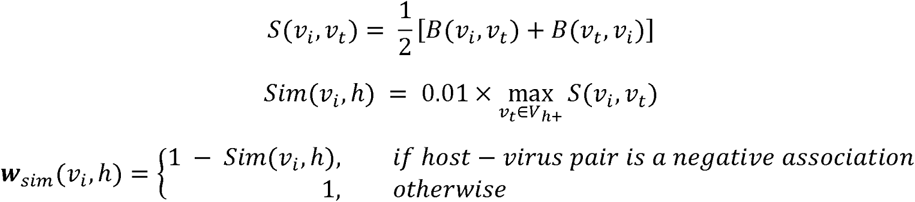

where *B*(*v_i,_ v_t_*) is the BLAST identity score between the virus *v_i_* (query) and virus *v_t_* (target), while *V_h_*_+_ represents viruses that have positive associations with host *h*. Furthermore, considering the imbalance between negative and positive pairs, we assigned a class weight ***w****_class_* to balance the impact of positive and negative associations.

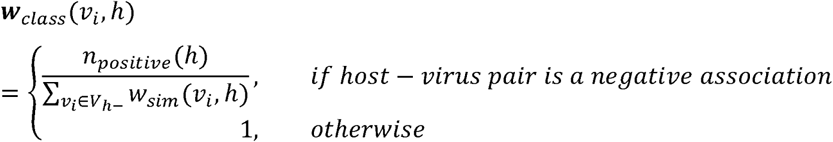

where *V_h_*_–_ is viruses having negative associations with host *h*. As our models were trained on subsequences generated from viral genomes, we also introduced a weight ***w****_length_* to reduce the impact of long viral genomes. We set the threshold for long genomes at 100,000 base pairs. Finally, we assigned a confidence weight ***w****_confidence_* for each host-virus pair during model training.

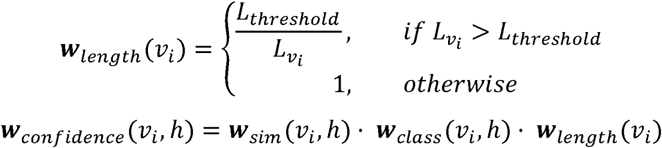

We scaled the contribution of each virus-host pair to the loss function using the confidence weight. Notably, confidence weights are only used during model training. For evaluation and prediction on new samples, confidence weights are all set to one.

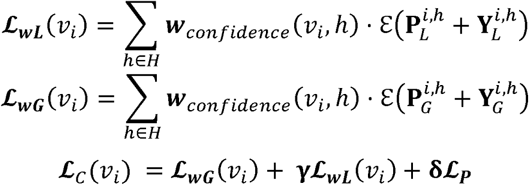

### VirHRanger predictive ensemble

#### Training

The six base models are trained separately on the five-fold cross-validation set, which includes genomic and protein foundation models, genomic and protein compositional models, a homology-based model, and a human-virus PPI-based model. A stacking strategy is applied^68^, employing linear regression as the meta-model. A separate meta-model is trained for each host category, resulting in a total of 79 meta-models in the VirHRanger predictive ensemble. The out-of-fold predicted probabilities of the base models during cross-validation are concatenated to form the input features for the meta-models.

#### Testing and prediction

When the predictive ensemble is benchmarked or applied to a new dataset, for each base model, the predictions from five models trained during cross-validation are averaged to calculate probabilities, which are then used as input for the meta-models.

### Viral sharing network analyses

We generated two unipartite viral sharing networks between animal hosts following the methodology^78^ of Carlson et al. Host order information was retrieved from the host-virus association dataset (with *Homo sapiens* separated from primates and used as the nodes of the different animal orders. These weights were calculated using the Jaccard index *J*(*A,B*), which network. The weights on the links between these nodes represent the shared viruses across is defined as:

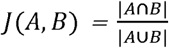

where *A* and *B* represent the number of representative viruses in two animal orders, |*A*∩*B*| is the number of shared representative viruses, and |*A*⋃*B*| is the total number of representative viruses across both orders. We separately created viral sharing networks for zoonotic and non-zoonotic viruses. The operational definition of a zoonotic virus was adapted from Olival et al.^3^, where a zoonotic virus is defined as any viral species detected in both humans and at least one other vertebrate host, without considering the direction of transmission. We also separately constructed viral sharing networks for documented and predicted host-virus associations. Finally, we visualized the networks by drawing bubble heatmaps using the *ggplot2* package^126^ in R with PhyloPic (http://www.phylopic.org).

### Predict for novel potential zoonotic and arboviruses

We downloaded all viral genomes and metadata from ZOVER database^80^ on June 19, 2022. A total of 51,016 genome sequences were retrieved. However, approximately 73.4% of these sequences were incomplete. To isolate viruses with complete genomes and remove highly duplicated records, we first extracted taxonomy information from the NCBI accession numbers using the *accessionToTaxa* function from the *taxonomizr* package^127^ in R. This process resulted in 4,438 viral species or strains. Together with the complete viral genomes downloaded from the NCBI Virus database, we obtained a total of 4,284 viral strains or species with complete genomes. To externally validate the performance of VirHRanger, we excluded the virus species that appeared in the training data. Metagenome-assembled genomes were removed due to concerns regarding their quality and taxonomic reliability. Additionally, for viruses of the same species, we used CD-HIT-EST to cluster viral genomes with a 90% identity threshold to reduce the influence of duplicated viral genomes. Finally, we obtained 315 representative viral genomes from 268 viral species that were not included in the training set of VirHRanger.

### Predict for coronaviruses

We downloaded all virus genomes from the NCBI virus database on November 10, 2022. We extracted a total of 1,810,864 coronaviruses with complete genomes. We retained all reference genomes from RefSeq and selected one representative genome for each virus species lacking a reference genome. This resulted in a total of 156 representative coronavirus genomes. The Maximum Likelihood phylogenetic tree was constructed using RAxML, version 8.2.12^128^, with a GTR model, 1,000 rapid bootstrap replications, and midpoint rooting method. Visualization and annotation of the tree were performed using the *ggtree* package^129^ in R, with branch lengths included for enhanced interpretability. For host range prediction, we excluded the genomes and host information of sarbecoviruses from the training data. We trained all models using five-fold cross-validation on the remaining genomes and performed a grid search to identify the best hyperparameter combination.

### Animal silhouettes used in figures

Animal silhouettes included in Figures 1, 2, 3, 4, and Supplementary Figures, which visually represent different animal orders, families, and species, were downloaded from PhyloPic (http://www.phylopic.org). All images are available for use under the Public Domain Dedication license. The silhouettes cover the following animal orders, families, and species:

- **Orders**: Anseriformes, Artiodactyla, Carnivora, Chiroptera, Diptera, Eulipotyphla, Galliformes, Ixodida, Lagomorpha, Lepidoptera, Passeriformes, Perissodactyla, Primates, and Rodentia
- **Families**: Anatidae, Bovidae, Camelidae, Canidae, Cercopithecidae, Cervidae, Cricetidae, Culicidae, Equidae, Felidae, Hominidae, Ixodidae, Leporidae, Muridae, Mustelidae, Phasianidae, Pteropodidae, Rhinolophidae, Sciuridae, Suidae, and Vespertilionidae
- **Species**: *Homo sapiens, Pan troglodytes, Chlorocebus aethiops, Macaca mulatta, Macaca fascicularis, Mus musculus, Rattus norvegicus, Bos taurus, Sus scrofa, Equus caballus, Capra hircus, Ovis aries, Canis lupus, Felis catus, Anas platyrhynchos,* and *Gallus gallus*

## Code availability

The trained individual models, ensemble models, and Python and Shell scripts for performing prediction are available for research purposes at https://github.com/JY-Bioinfo/VirHRanger.

## Data availability

The genomic and protein sequences for pretraining and fine-tuning were obtained from publicly available resources. The pretraining genomic sequences were obtained from https://viralzone.expasy.org/8676. The genomic and protein sequences for fine-tuning were obtained from https://ftp.ncbi.nlm.nih.gov/genomes/refseq/ and https://ftp.ncbi.nlm.nih.gov/genbank/. The original animal-virus association records were obtained from https://www.viralemergence.org/virion, https://www.genome.jp/virushostdb/, https://www.cdc.gov/arbocat/, and http://v2.insect-genome.com/. The human virus PPI information was acquired from http://zzdlab.com/hvidb/. The novel zoonotic and arboviruses were obtained from https://www.mgc.ac.cn/cgi-bin/ZOVER/main.cgi.

## Acknowledgments

This work was supported by the National Key Research and Development Program of China (2021YFC2300300) and the National Natural Science Foundation of China (32070667, 31671366, 32300078, T2321001). Part of the analysis was performed on the High-Performance Computing Platform of Peking University and the Biomedical Computing Platform of the National Biomedical Imaging Center of Peking University.

## Author contributions

H.Q.Z. and X.Q.J. co-supervised the study. J.Y.G. and Q.G. carried out the algorithm design. J.Y.G., Q.G., H.C.Y., P.X.G., J.H.H. and H.Y.Z. performed the analysis. Y.L.H., J.T., M.L., and Y.Z. provided advice on study design and analyses. J.Y.G., Q.G., H.Q.Z., and X.Q.J. wrote and revised the paper with input from all co-authors. All the authors read and accepted the final version of the manuscript.

