## Supplementary Material for "Developing Foundation Models for Predicting Viral Animal Host Range in Intelligent Surveillance"

**This PDF file includes:**

Supplementary Figure S1 to S15

**Other Supplementary Material for this manuscript includes the following:**

Supplementary Table 1 (.xlsx): A comprehensive collection of animal-virus associations

Supplementary Table 2 (.xlsx): Representative viral genomes and animal hosts

Supplementary Table 3 (.xlsx): Performance of VirHRanger across 79 host categories

Supplementary Table 4 (.xlsx): Performance of VirHRanger across different host taxonomic ranks by macro-averaging

Supplementary Table 5 (.xlsx): Performance of all models for human infecting viruses

Supplementary Table 6 (.xlsx): Sample-wise host range prediction performance of VirHRanger and Homology-based model

Supplementary Table 7 (.xlsx): Identification of known zoonotic viruses and prediction of additional zoonotic viruses by VirHRanger

Supplementary Table 8 (.xlsx): Identification of known arboviruses and prediction of additional arboviruses by VirHRanger

Supplementary Table 9 (.xlsx): External validation of VirHRanger on potential zoonotic and arboviruses

Supplementary Table 10 (.xlsx): Host range predictions for coronaviruses by VirHRanger

Supplementary Table 11 (.xlsx): AUROC values from 5-fold cross-validation for models trained on human-virus PPIs and SLiMs

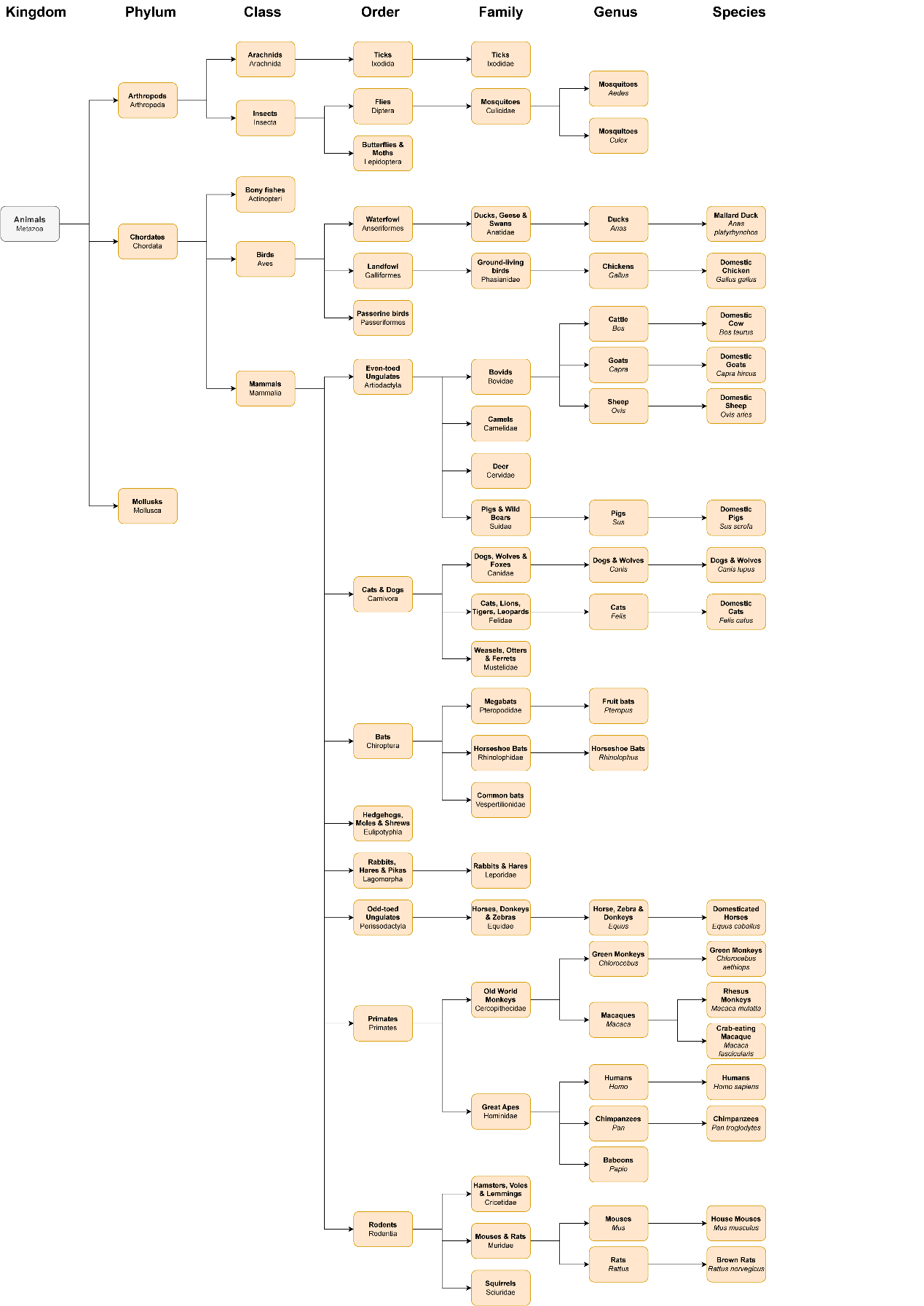

**Supplementary Figure S1. Hierarchy of host categories predicted by VirHRanger.** Seventy-nine host categories spanning different taxonomic levels were retrained to allow VirHRanger to systematically predict animal host range for viruses. The common name and scientific name of each host category are depicted in rounded rectangles. Figures were created using diagrams.net (https://www.diagrams.net).

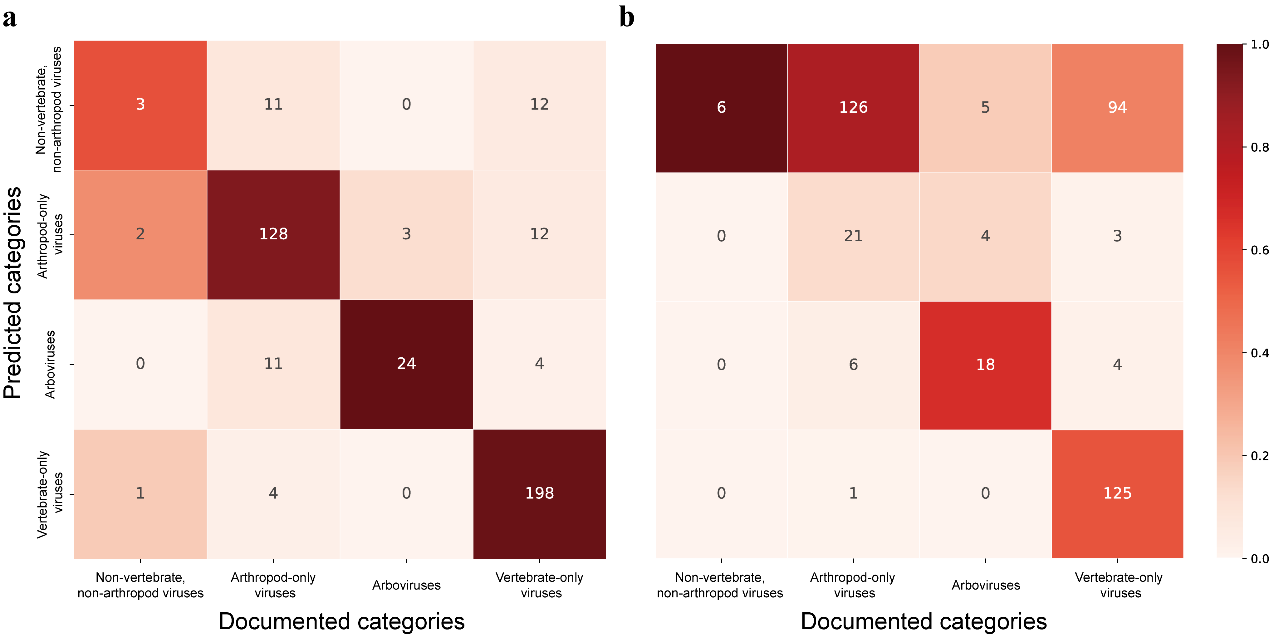

**Supplementary Figure S2. Performance of VirHRanger and the homology-based model when applied to arthropod-associated viruses.** The heatmaps compare documented categories and predicted categories of viruses based on host specificity, including non-vertebrate, non-arthropod viruses, arboviruses, arthropod-only viruses, and vertebrate-only viruses. The prediction models are **(a)** VirHRanger and **(b)** the homology-based model.

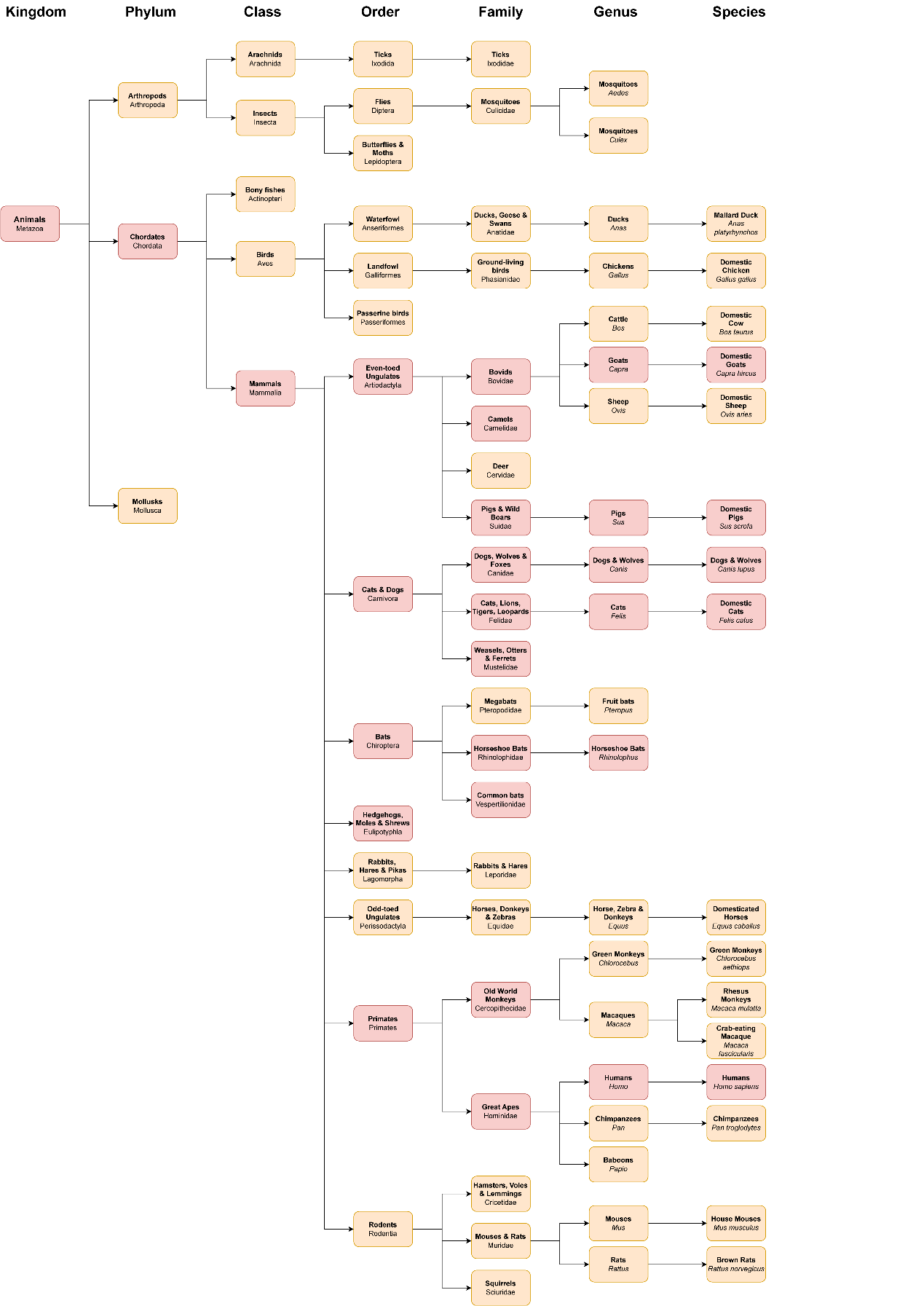

**Supplementary Figure S3. Animal host range of SARS-CoV-2 predicted by VirHRanger.** According to the prediction of VirHRanger, animal host categories associated with SARS-CoV-2 are colored in red. Figures were created using diagrams.net (https://www.diagrams.net).

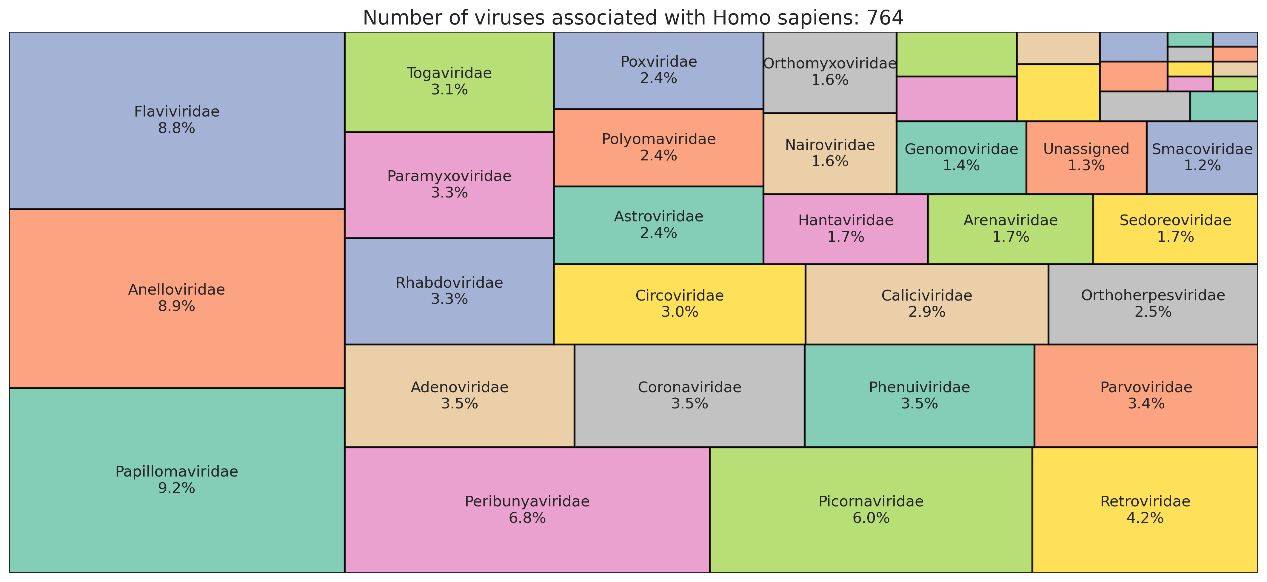

**Supplementary Figure S4. The composition of viruses associated with humans.** The treemap illustrates the diversity of viral families associated with humans in the training dataset of VirHRanger. Each segment represents a viral family, labeled with its scientific name and relative abundance. Viral families with an abundance of less than 1% are excluded for readability.

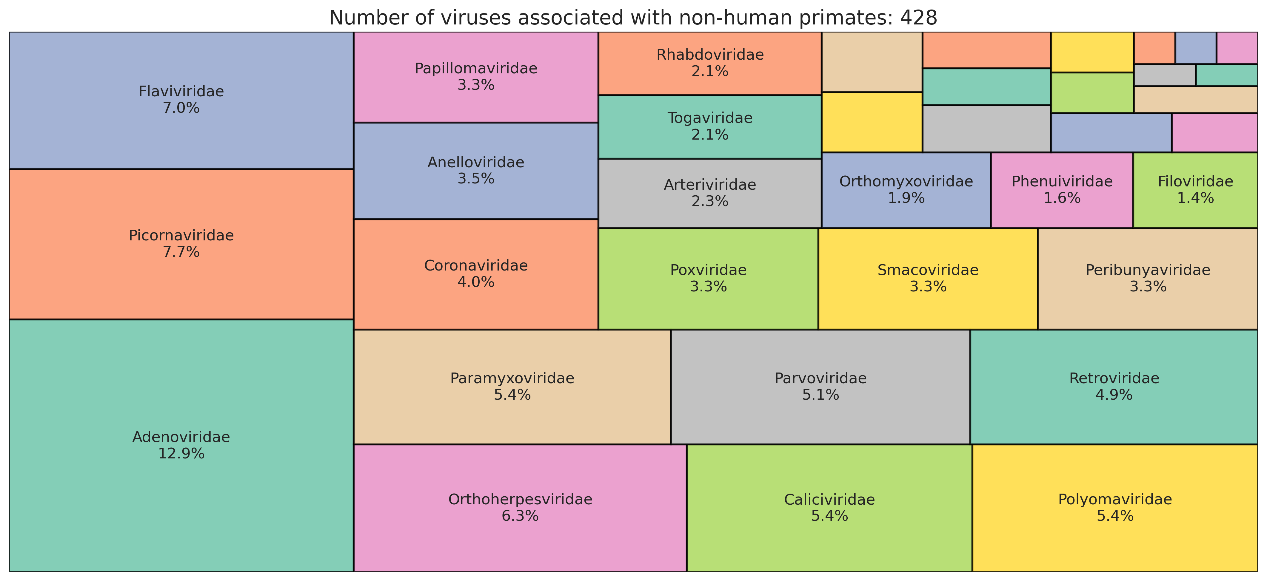

**Supplementary Figure S5. The composition of viruses associated with non-human primates.** The treemap illustrates the diversity of viral families associated with non-human primates in the training dataset of VirHRanger. Each segment represents a viral family, labeled with its scientific name and relative abundance. Viral families with an abundance of less than 1% are excluded for readability.

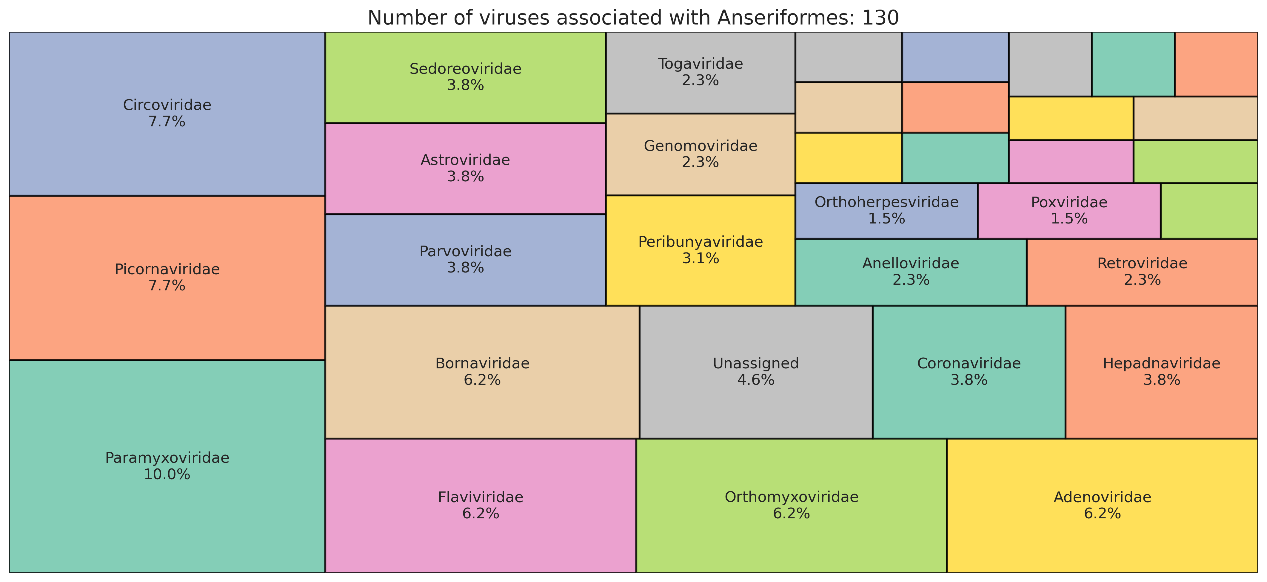

**Supplementary Figure S6. The composition of viruses associated with Anseriformes.** The treemap illustrates the diversity of viral families associated with Anseriformes in the training dataset of VirHRanger. Each segment represents a viral family, labeled with its scientific name and relative abundance. Viral families with an abundance of less than 1% are excluded for readability.

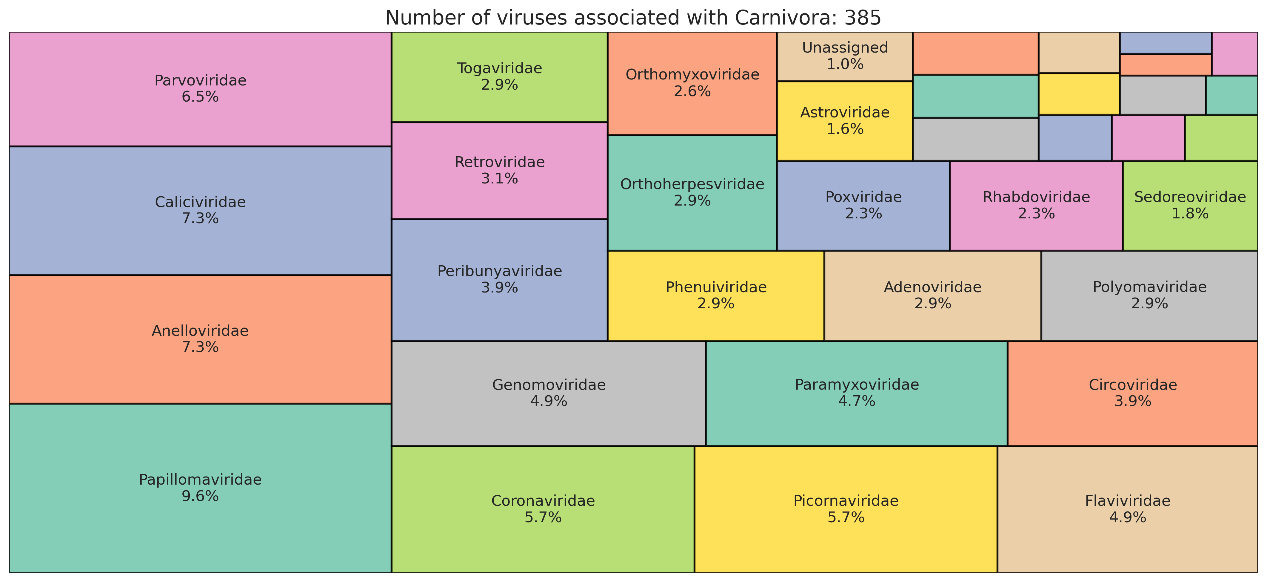

**Supplementary Figure S7. The composition of viruses associated with Carnivora.** The treemap illustrates the diversity of viral families associated with Carnivora in the training dataset of VirHRanger. Each segment represents a viral family, labeled with its scientific name and relative abundance. Viral families with an abundance of less than 1% are excluded for readability.

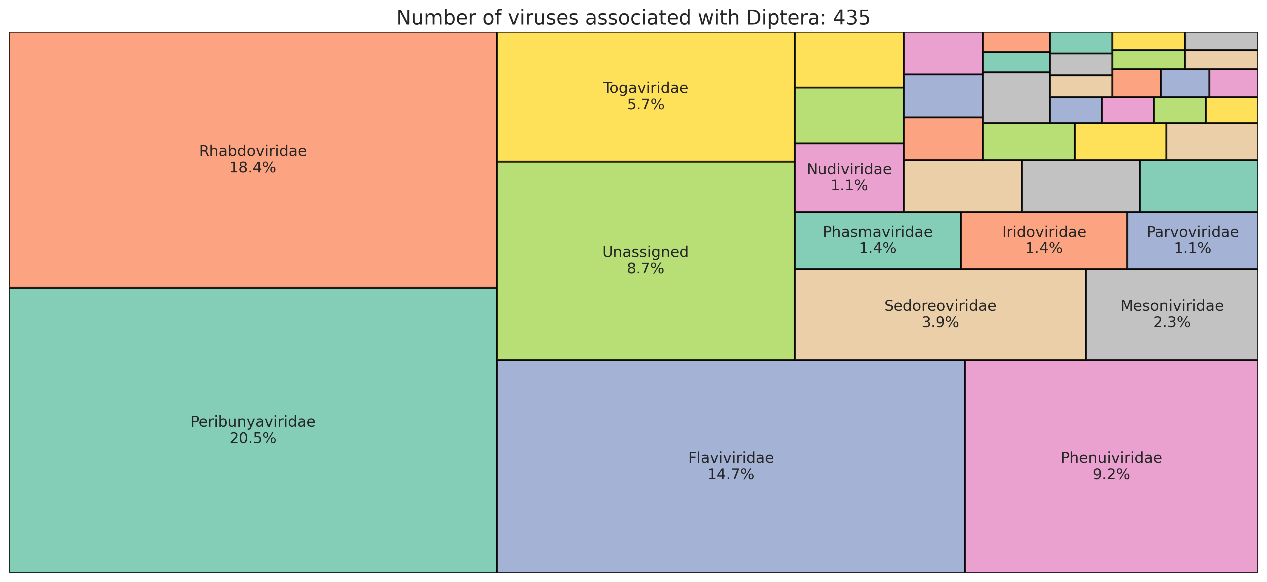

**Supplementary Figure S8. The composition of viruses associated with Diptera.** The treemap illustrates the diversity of viral families associated with Diptera in the training dataset of VirHRanger. Each segment represents a viral family, labeled with its scientific name and relative abundance. Viral families with an abundance of less than 1% are excluded for readability.

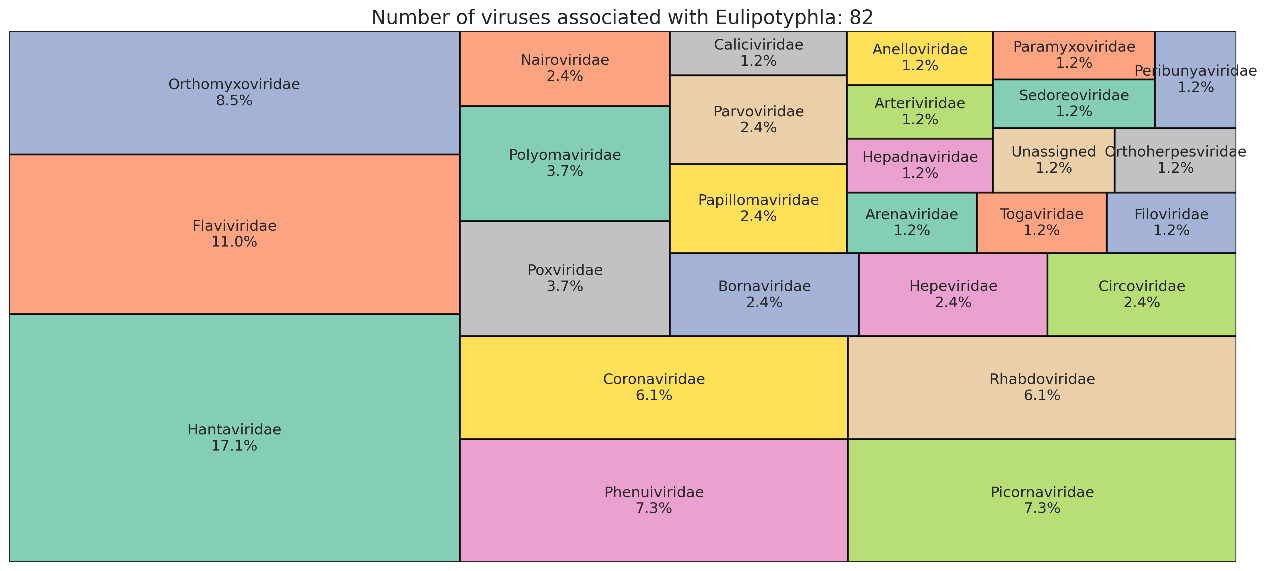

**Supplementary Figure S9. The composition of viruses associated with Eulipotyphla.** The treemap illustrates the diversity of viral families associated with Eulipotyphla in the training dataset of VirHRanger. Each segment represents a viral family, labeled with its scientific name and relative abundance. Viral families with an abundance of less than 1% are excluded for readability.

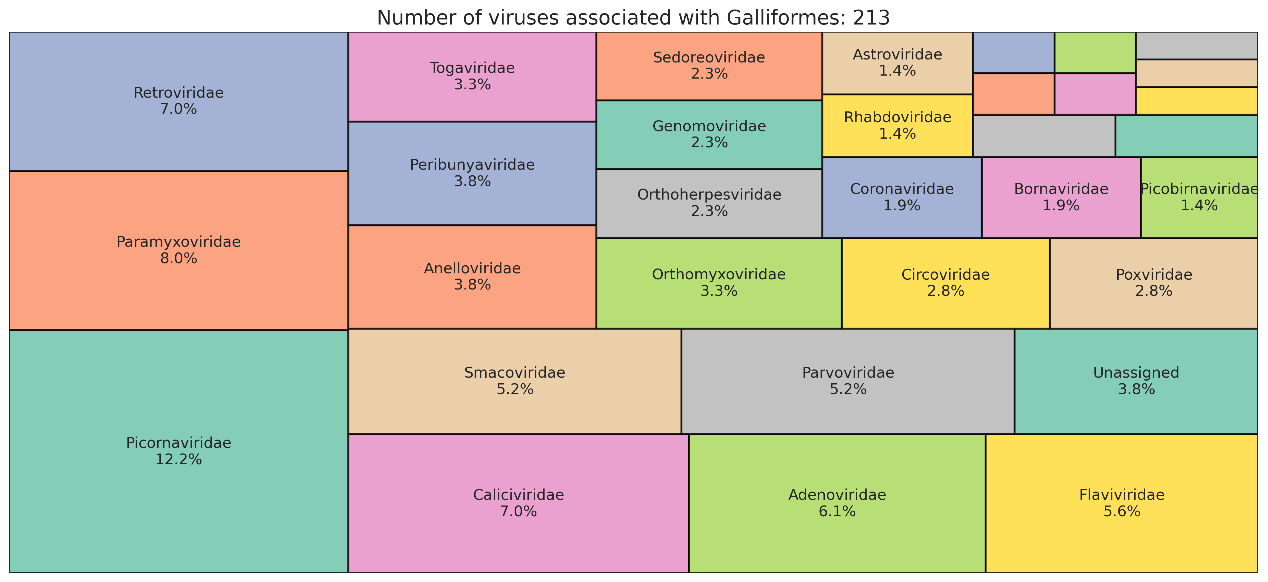

**Supplementary Figure S10. The composition of viruses associated with Galliformes.** The treemap illustrates the diversity of viral families associated with Galliformes in the training dataset of VirHRanger. Each segment represents a viral family, labeled with its scientific name and relative abundance. Viral families with an abundance of less than 1% are excluded for readability.

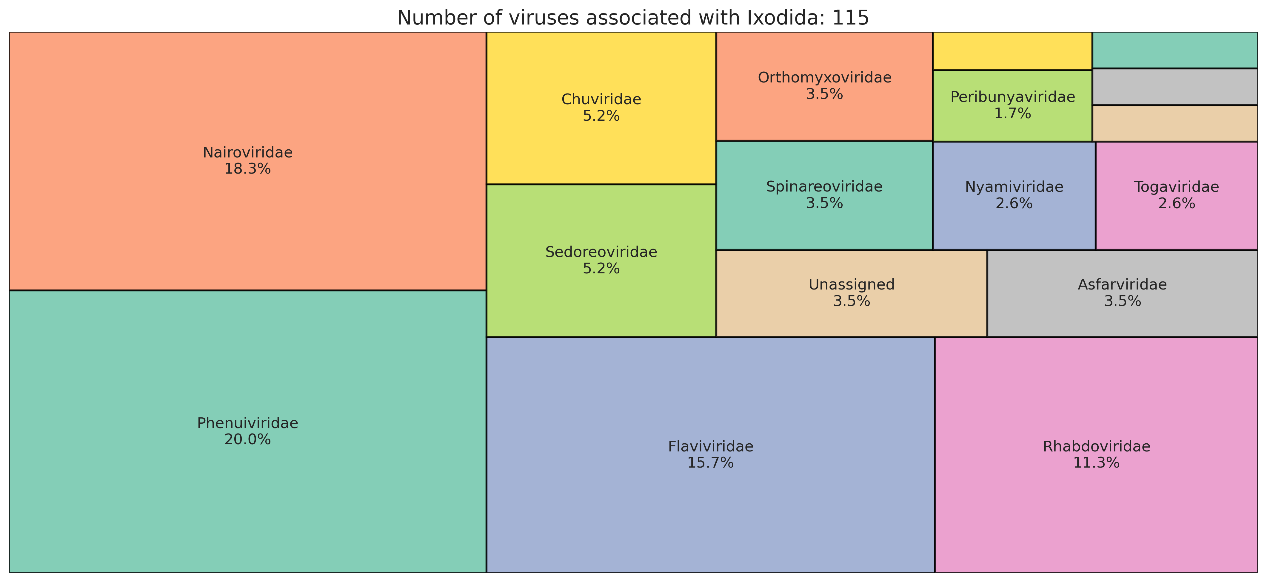

**Supplementary Figure S11. The composition of viruses associated with Ixodida.** The treemap illustrates the diversity of viral families associated with Ixodida in the training dataset of VirHRanger. Each segment represents a viral family, labeled with its scientific name and relative abundance. Viral families with an abundance of less than 1% are excluded for readability.

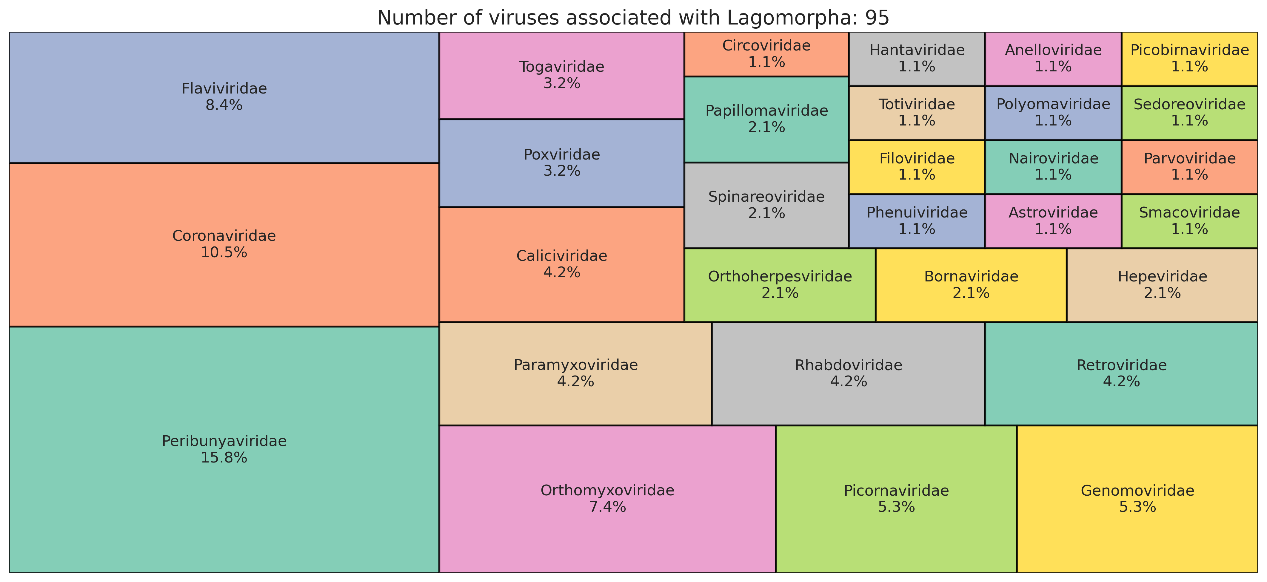

**Supplementary Figure S12. The composition of viruses associated with Lagomorpha.** The treemap illustrates the diversity of viral families associated with Lagoporpha in the training dataset of VirHRanger. Each segment represents a viral family, labeled with its scientific name and relative abundance. Viral families with an abundance of less than 1% are excluded for readability.

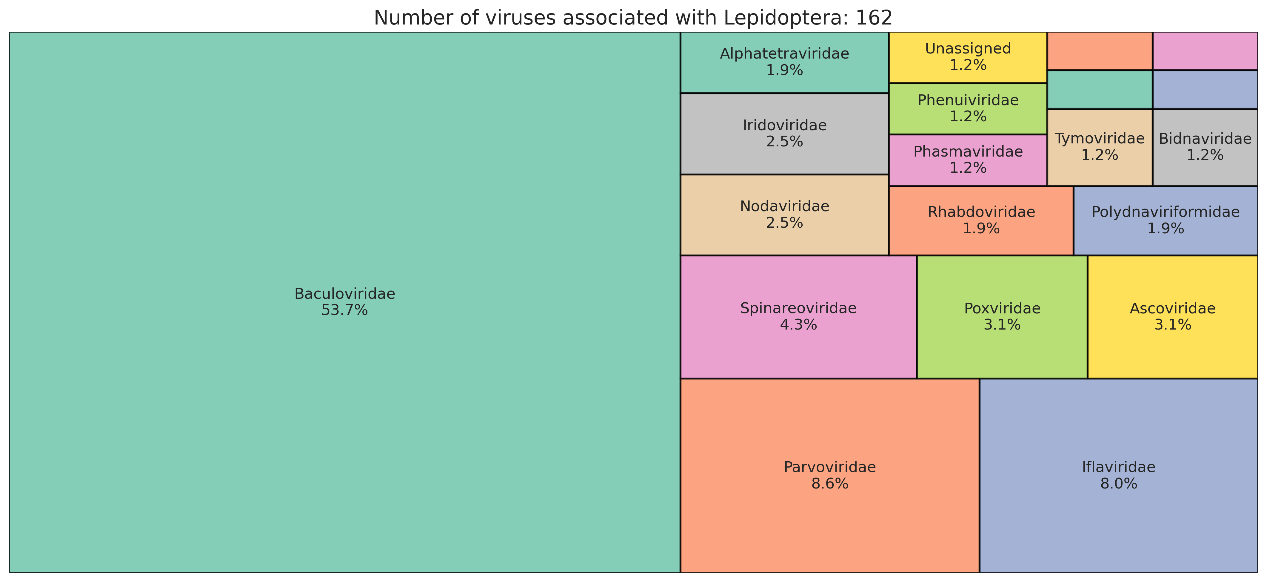

**Supplementary Figure S13. The composition of viruses associated with Lepidoptera.** The treemap illustrates the diversity of viral families associated with Lepidoptera in the training dataset of VirHRanger. Each segment represents a viral family, labeled with its scientific name and relative abundance. Viral families with an abundance of less than 1% are excluded for readability.

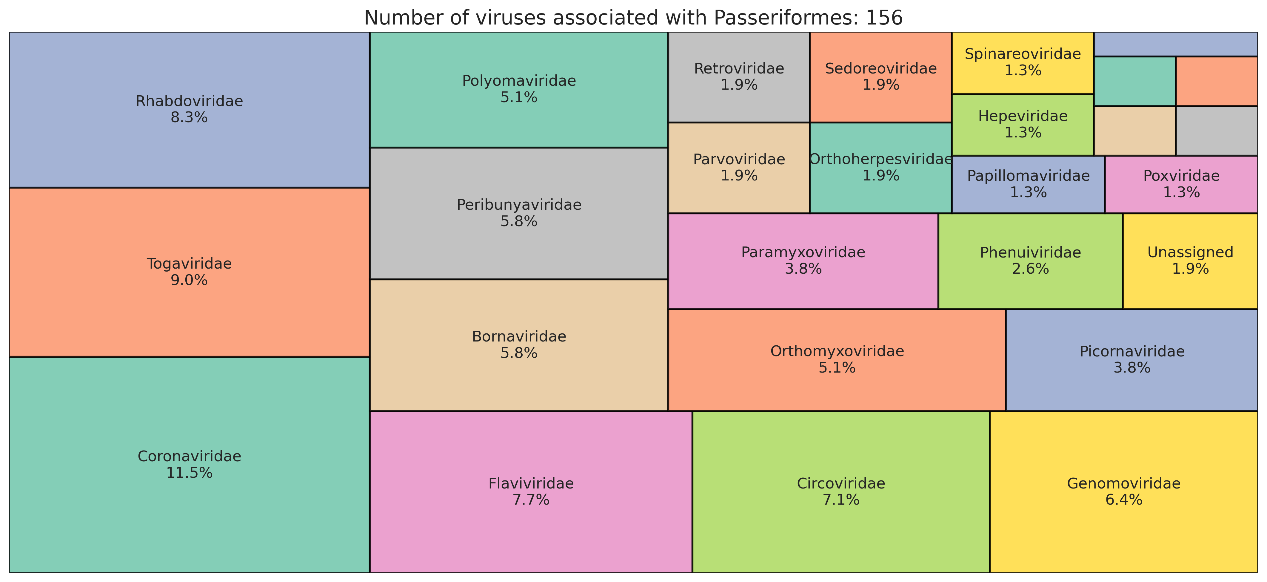

**Supplementary Figure S14. The composition of viruses associated with Passeriformes.** The treemap illustrates the diversity of viral families associated with Passeriformes in the training dataset of VirHRanger. Each segment represents a viral family, labeled with its scientific name and relative abundance. Viral families with an abundance of less than 1% are excluded for readability.

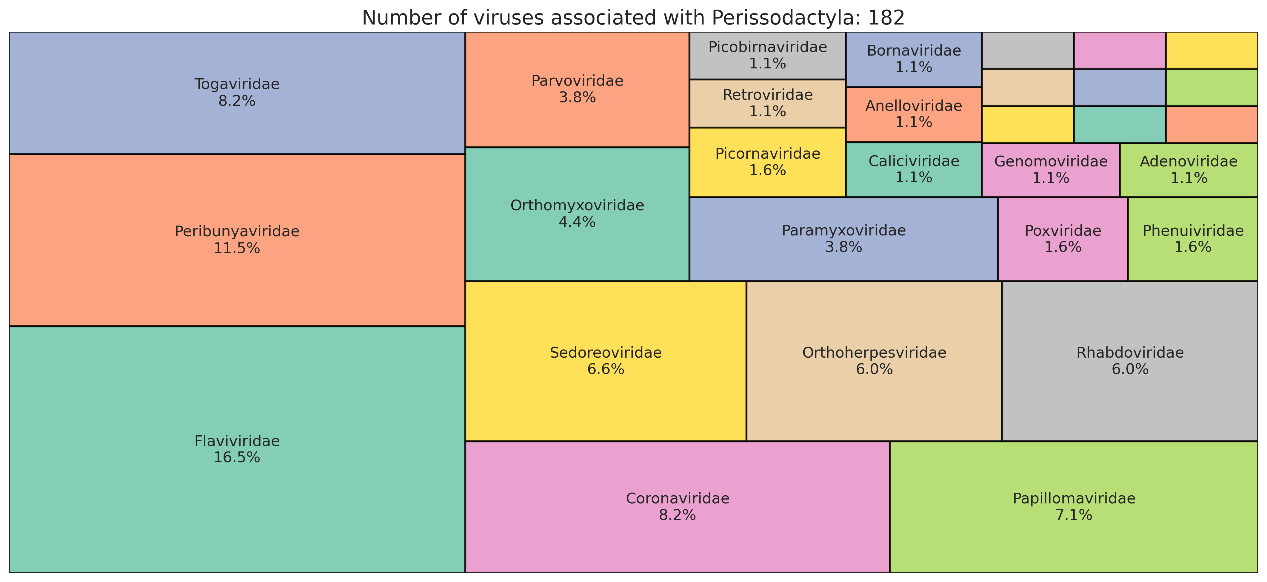

**Supplementary Figure S15. The composition of viruses associated with Perissodactyla.** The treemap illustrates the diversity of viral families associated with Perissodactyla in the training dataset of VirHRanger. Each segment represents a viral family, labeled with its scientific name and relative abundance. Viral families with an abundance of less than 1% are excluded for readability.
